# *MutS-Homolog2* silencing generates tetraploid meiocytes in tomato (*Solanum lycopersicum*)

**DOI:** 10.1101/142612

**Authors:** Supriya Sarma, Arun Kumar Pandey, Maruthachalam Ravi, Yellamaraju Sreelakshmi, Rameshwar Sharma

## Abstract

MSH2 is the core protein of MutS-homolog family involved in recognition and repair of the errors in the DNA. While other members of MutS-homolog family reportedly regulate mitochondrial stability, meiosis, and fertility, MSH2 is believed to participate mainly in mismatch repair. The search for polymorphism in *MSH2* sequence in tomato accessions revealed both synonymous and nonsynonymous SNPs; however, SIFT algorithm predicted that none of the SNPs influenced MSH2 protein function. The silencing of *MSH2* gene expression by RNAi led to phenotypic abnormalities in highly-silenced lines, particularly in the stamens with highly reduced pollen formation. *MSH2* silencing exacerbated formation of UV-B induced thymine dimers and blocked light-induced repair of the dimers. The *MSH2* silencing also affected the progression of male meiosis to a varying degree with either halt of meiosis at zygotene stage or formation of diploid tetrads. The immunostaining of male meiocytes with centromere localized CENPC (Centromere protein C) protein antibody showed the presence of 48 univalent along with 24 bivalent chromosomes suggesting abnormal tetraploid meiosis. The mitotic cells of root tips of silenced lines showed diploid nuclei but lacked intervening cell plates leading to cells with syncytial nuclei. Thus we speculate that tetraploid pollen mother cells may have arisen due to the fusion of syncytial nuclei before the onset of meiosis. It is likely that in addition to Mismatch repair (MMR), MSH2 may have an additional role in regulating ploidy stability.

## INTRODUCTION

The survival of the living organisms is dependent on the precise and error-free replication of DNA. In addition, normal metabolic activity and environmental factors such as radiation also cause damage to the DNA. The fidelity of the DNA is ensured by a cohort of mechanisms that detects and eliminates mistakes occurring during DNA replication. At least five major repair pathways are known to operate in higher plants: base excision repair, nucleotide excision repair, mismatch repair (MMR) and doublestrand break repair comprising of homologous recombination repair and nonhomologous end joining (Takata *et al*., 1998; Guarné *et al*., 2004; Reardon and Sancar, 2005; Jacobs and Schär, 2012). Among these, MMR is a major pathway that corrects base–base and insertion–deletion mismatches generated during DNA replication or mutagenesis (Dowen *et al*., 2010; Srivatsan *et al*., 2014).

MMR is a highly conserved pathway that exists in all living organisms. The MMR process consists of three key steps: mismatch recognition, excision, and resynthesis (Ban *et al*., 1999; Antony and Hingorani, 2003; Guarné *et al*., 2004). The first step of the mismatch recognition in eukaryotes is carried out by the homologs of prokaryotic MutS proteins, namely MSH protein subunits. There are eight homologs of MutS in eukaryotes: MSH1 to MSH8, of which MSH7 is found only in plants (Culligan and Hays, 2000) and MSH8 in Euglenozoa (Sachadyn, 2010). The MSH proteins recognize mismatches as heterodimers, where MutSα (MSH2-MSH6) repairs base-base mismatches or 1-2 nucleotide insertion-deletion loops (Acharya *et al*., 1996; Genschel *et al*., 1998), while MutSβ (MSH2-MSH3) recognizes larger, up to 14 nucleotide insertion-deletion loops (Modrich, 1991; Marti *et al*., 2002). Plants form an additional heterodimeric complex known as MutSγ (MSH2–MSH7) (Culligan and Hays, 2000) which recognizes some base–base mismatches and reportedly plays a role in meiotic recombination (Lloyd *et al*., 2007). The binding of MutSα/β to mismatched DNA initiates conformational changes in MutS recruiting MutL complex followed by activation of exonuclease1/PCNA. The gap generated by DNA excision is filled by action of DNA polymerase and DNA ligase.

It is reported that the other members of MSH2 family function beyond MMR. MSH1 is required for mitochondrial stability (Reenan and Kolodner, 1992) and disruption of MSH1 can influence male sterility in several species (Sandhu *et al*., 2007; Zhao *et al*., 2016). The MSH4 and MSH5 form heterodimer between them without involving MSH2 and exclusively function in meiosis (Hollingsworth *et al*., 1995; Schofield and Hsieh, 2003, Lu *et al*., 2008; Wang *et al*., 2016). The expression of MSH7 is required for wild-type level of fertility in barley (Lloyd *et al*., 2007). MSH7 also plays a role in UV-B-induced DNA damage recognition and recombination in Arabidopsis (Lario *et al*., 2015).

Among the MMR proteins, MSH2 is a common protein in three of the heterodimers formed in the plants. In conformity of the role of MSH2 protein in repair of replication errors, an Arabidopsis mutant defective in MSH2 shows microsatellite instability (Depeiges *et al*., 2005). The *msh2* mutants in Moss show strong developmental defects, are sterile and have a mutator phenotype (Trouiller *et al*., 2006). A similar phenotype was also observed in *Arabidopsis thaliana msh2* mutant lines albeit only after five generations (Hoffman *et al*., 2004). Considering that suppression of MMR leads to an increase in natural mutation frequency, *MSH2* gene was silenced in *Nicotiana tabacum* and *N. plumbaginifolia* using an artificial microRNA (amiRNA). Though the transformed lines showed developmental and morphological abnormality, the mutant phenotype was not transmitted to subsequent generations (van Marcke and Angenon, 2013).

Among the interacting partners for MSH2; mutants are reported only for *MSH6* and *MSH7* in Arabidopsis. The *msh6* mutant does not display a visible phenotype (Lario *et al*., 2011) however, *msh7* mutant shows an increase in meiotic recombination frequency (Lario *et al*., 2015). The role of *MSH7* in meiotic recombination is also reported in tomato, where RNAi silencing of *MSH7* improves homeologous meiotic recombination with a chromosome from *S. lycopersicoides* (Tam *et al*., 2011). Most likely the high increase in meiotic recombination is mediated by the MSH2-MSH7 complex as Arabidopsis *msh2* mutant also shows 40% increase in the meiotic crossover rates (Emmanuel *et al*., 2006).

In this study, our aim was to isolate MSH2-defective mutants to obtain a hypermutable line to increase the efficiency of EMS-mutagenesis in tomato. We screened natural accessions of tomato and EMS-mutagenized lines for SNPs/mutation in tomato *MSH2* by EcoTILLING and TILLING respectively. Considering that little polymorphism was observed in *MSH2* in tomato accessions, we silenced *MSH2* gene using *MSH2*-RNAi construct. We report that silencing of *MSH2* gene led to a developmental defect in tomato lines with abnormal flowers bearing tetraploid meiocytes in the anthers. The examination of root cells showed the formation of syncytial nuclei and defects in the cell plate formation. Our study indicates that *MSH2* may also have a role in the regulation of meiotic and mitotic cell divisions.

## RESULTS

### EcoTILLING of *MSH2* gene showed little polymorphism in tomato accessions

Given the pivotal role of *MSH2* in MMR, we screened a total of 391 tomato accessions for polymorphism in the gene. A 2.7 Kb region (4-9 exon) of *MSH2* predicted to be deleterious by CODDLE software was chosen for EcoTILLING. Consistent with the essential nature of the gene, only 7 accessions showed polymorphism with a total of 13 SNPs with a frequency of 1.23 SNP/100 Kb. (Figure S1). Among the identified SNPs, 6 were non-synonymous, and 7 were synonymous (Table 1). The likelihood of non-synonymous SNPs affecting protein function was examined by SIFT software. Since none of the SNP showed a SIFT score <0.05, these SNPs were predicted to be non-deleterious in nature. A parallel screening of 1243 bp region of *MSH2* gene covering 4th exon of 11304 tomato EMS-mutagenized lines by TILLING did not yield any mutation.

**Table 1.** List of SNPs detected in *MSH2* gene in different tomato accessions along with amino acid changes. The scores predicted by SIFT suggests that none of the amino acid changes would affect MSH2 protein activity. **Note:** A SIFT score of less than 0.05 predicts the base substitution to be intolerant.

| Accession No. | No. of SNPs | Base Change | Amino acid change | SIFT Score | Nature of SNP |
| --- | --- | --- | --- | --- | --- |
| EC398695 | 2 | G1168T | A77A | - | Synonymous |
|  |  | T1365C | V143A | 1.00 | Nonsynonymous |
| EC520075 | 3 | A1720T | A261A | - | Synonymous |
|  |  | T1876G | A313A | - | Synonymous |
|  |  | G3193T | V639V | - | Synonymous |
| EC520052 | 1 | T1876G | A313A | - | Synonymous |
| EC521078 | 4 | C1312T | F125F | - | Synonymous |
|  |  | G1545T | G203V | 0.23 | Nonsynonymous |
|  |  | G1808A | D291N | 0.83 | Nonsynonymous |
|  |  | G2733T | G545V | - | Nonsynonymous |
| EC528374 | 3 | G1168T | A77A | - | Synonymous |
|  |  | T1365C | V143A | 1.00 | Nonsynonymous |
|  |  | T1876G | A313A | - | Synonymous |
| EC34480 | 4 | C1312T | F125F | - | Synonymous |
|  |  | G1381C | A148A | - | Synonymous |
|  |  | G1808A | D291N | 0.83 | Nonsynonymous |
|  |  | G3777T | A707V | - | Nonsynonymous |
| EC520078 | 4 | C1312T | F125F | - | Synonymous |
|  |  | T1339C | A134A | - | Synonymous |
|  |  | C1796G | Q287E | 0.37 | Nonsynonymous |
|  |  | G1808A | D291N | 0.83 | Nonsynonymous |

### Generation of *MSH2*-RNAi lines

Considering recalcitrance to the mutagenesis and low polymorphism in *MSH2* gene, we silenced the gene by RNAi. A 301 bp region of *MSH2* gene with no significant homology to any other tomato gene was amplified and cloned into pHELLSGATE 12 vector (Figure S2). A total of ten independent transgenic lines bearing *MSH2*-RNAi construct were raised using *Agrobacterium*-mediated transformation. Southern blot analysis of T_0_ lines revealed the presence of the transgene in five lines (Figure S3). In T_2_ generation out of three lines, the lines 1T_2_1-11 and 2T_2_7-5 with strong MSH2 silencing showed a single transgene copy (Figure S4).

### MMR transcript analysis in *MSH2*-RNAi lines using real-time PCR

To ascertain the magnitude of gene silencing, the level of *MSH2* transcript in transgenic lines was quantified by real-time PCR. Out of five Southern positive lines, the line 1T_2_1-11 showed ~80% reduction of *MSH2* transcript level followed by line 2T_2_5-5 and 2T_2_7-5 compared to wild-type (WT) control (Figure 1). Line 3T_2_1-4 showed moderate, and 5T_2_3-3 showed no reduction in *MSH2* transcript levels. The reduction in the transcript level in line1T21-11 was not restricted to *MSH2* alone; it also reduced the transcript levels of *MSH5* and *MSH7* albeit to a moderate but a significant extent. Since, *MSH2, MSH5*, and *MSH7* genes bear no significant sequence homology, the reduction in *MSH5* and *MSH7* transcript is most probably due to some indirect influence of *MSH2* transcript reduction.

**Figure 1.**
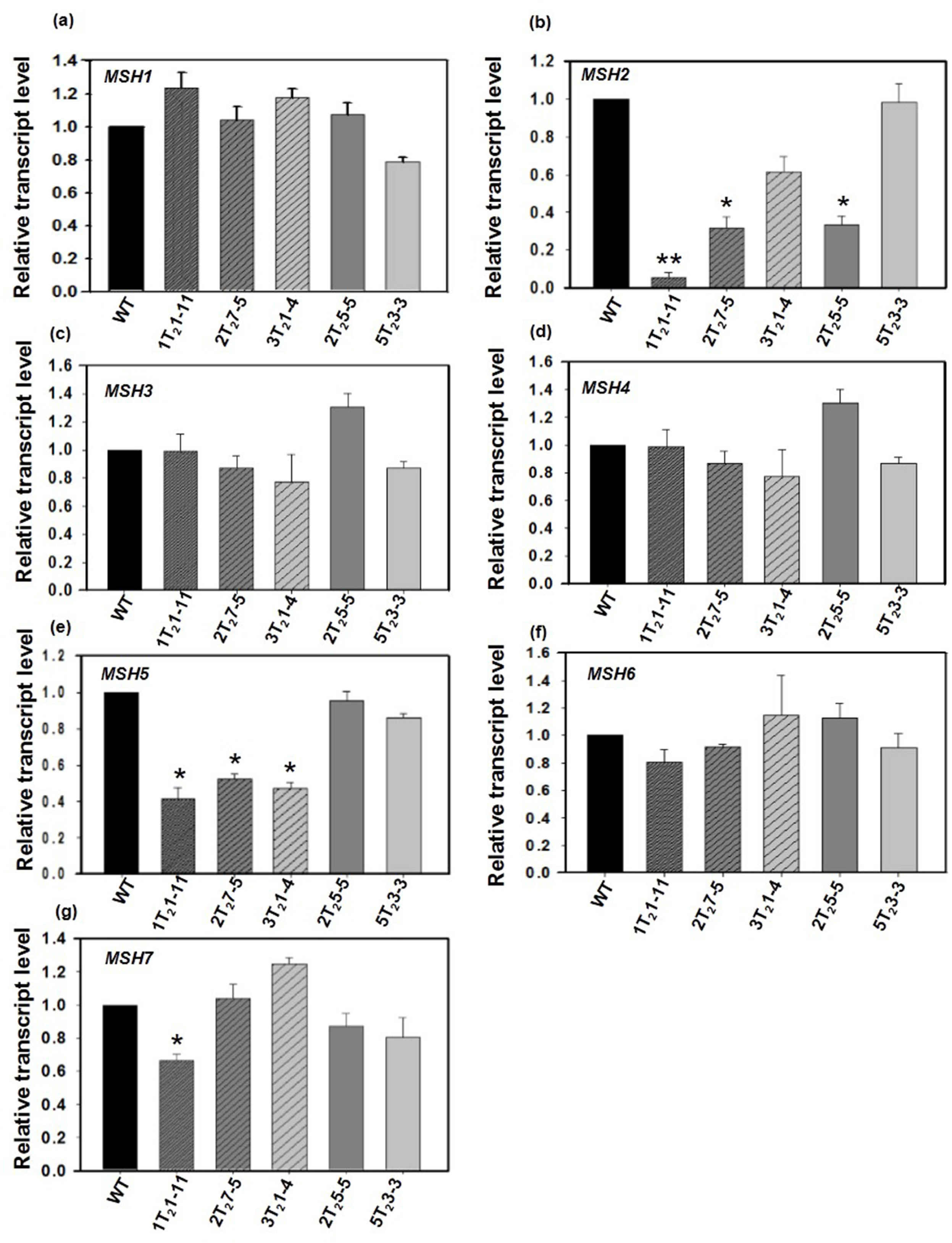
Relative transcript levels of members of *MSH* gene family in leaves of five different *MSH2*-RNAi lines. (a) *MSH1*, (b) *MSH2*, (c) *MSH3*, (d) *MSH4*, (e) *MSH5*, (f) *MSH6*, (g) *MSH7*. The transcripts levels were expressed after normalization with two internal controls, *β-actin* and *ubiquitin*. Note: line No. 1T_2_1-11 shows the reduction in *MSH2, MSH5*, and *MSH7* transcript. The statistical significance was determined by Student’s t-test. * P<0.05.

### *MSH2* silencing exacerbates UV-B-mediated thymine dimer formation

In mammalian cells, the contribution of the MMR pathway to the UVB-induced DNA damage is well established (Narine *et al*., 2007; Seifert *et al*., 2008). Similarly, Arabidopsis *MSH2* T-DNA insertion lines show higher levels of Cyclobutane Pyrimidine Dimer (CPD) on exposure to UV-B (Lario *et al*., 2011, 2015). We used UV-B induced thymine dimer formation as an indirect assay to monitor the activity of MSH2 in transgenic plants in comparison with WT. Pre- and post-UV-B exposure, the thymine dimers were quantified in genomic DNA of above plants by using a thymine dimer-specific monoclonal antibody (Figure S5, Figure 2). Prior to UV-B exposure, the steady-state levels of thymine dimers in control and *MSH2* silenced lines were nearly similar. However, post-6 h UV-B exposure, *MSH2* silenced lines showed a high level of dimers than the control plants. The thymine dimer level in *MSH2*-RNAi lines was higher than the control even after 24 h dark or white light incubation (Figure 2). It appears that high accumulation of thymine dimer in M*SH2* transgenic lines may have resulted from low *in-vivo* MMR activity.

**Figure 2.**
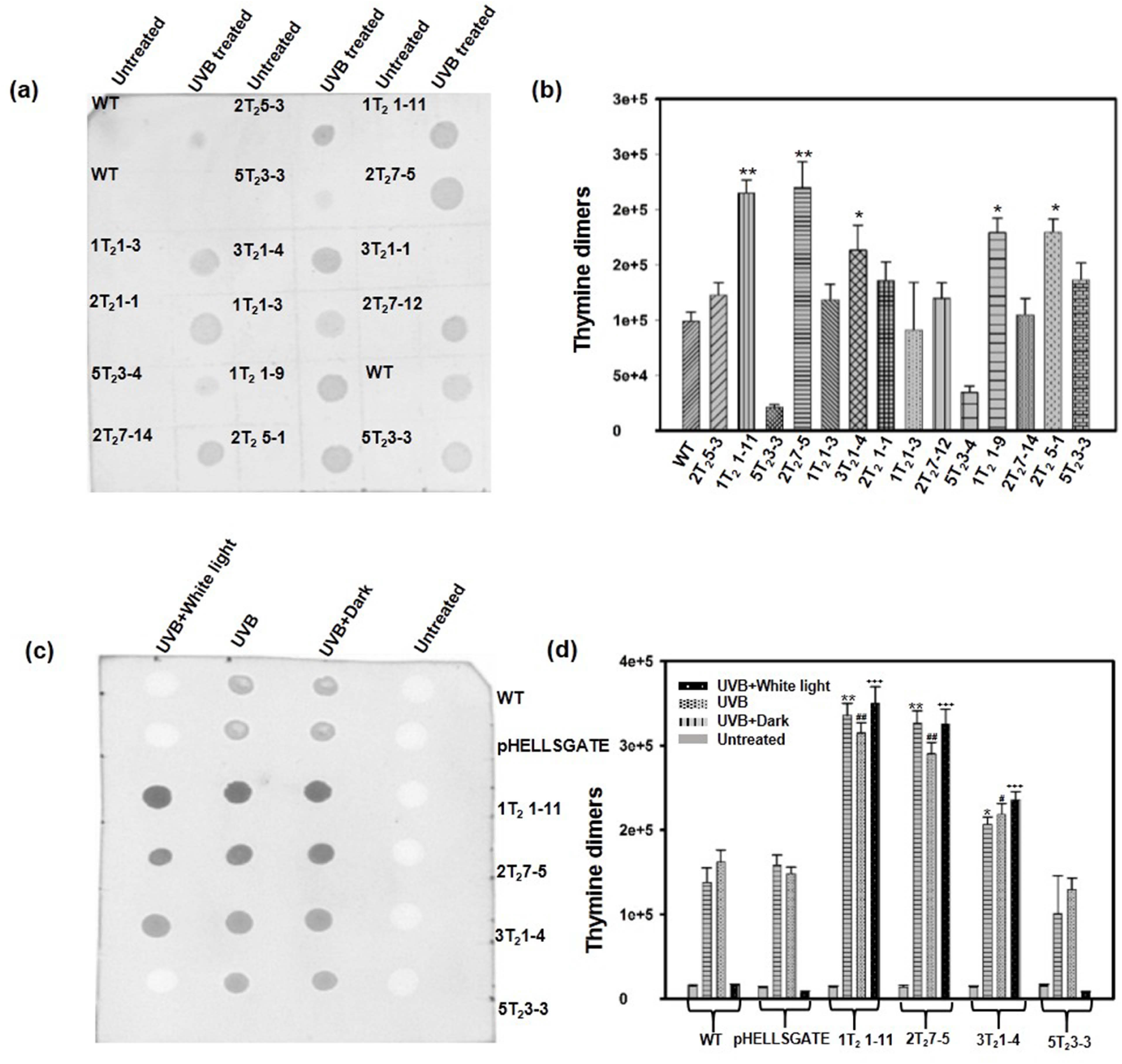
Determination of the *in-vivo* activity of MSH2 by quantification of thymine dimer levels using a thymine dimer-specific antibody in UV-B irradiated WT and MSH2-RNAi lines by dot-blot assay. (a, b) Quantification of thymine dimers in WT and *MSH2*-RNAi plants after 6 h with or without UV-B exposure (untreated). (a) Representative dot-blot. The label on top of each spot indicates the line number of *MSH2* silenced line. (b) Quantification of thymine dimers in WT and *MSH2*-RNAi lines by Image J analysis. (c, d) Quantification of thymine dimers in WT and *MSH2*-RNAi plants. Thymine dimers were quantified either immediately after 6 h exposure with UV-B, or after 24 h of white light, or dark incubation. (c) Representative dot-blot. The labels on the right of the blot indicate the line number of *MSH2* silenced line. (d) Quantification of thymine dimers in *MSH2*-RNAi lines by Image J analysis. Note: The thymine dimers in (b) and (d) were quantified using three independent biological replicates. The statistical significance was determined by Student’s t-test. * P<0.05.

### *MSH2*-RNAi lines display abnormal pollen formation

The lines 1T_2_1-11 and 2T_2_7-5 with a higher reduction in *MSH2* transcript also showed phenotypic abnormalities in T_1_ and T_2_ generation (Figure 3a-f). The most pronounced abnormality was in the stamen morphology (Figure 3g-h). The plants in later generations too showed abnormalities in flowers, mostly in the stamens. The progeny plants of silenced lines either failed to set fruits or set fruits with very few seeds (Figure 3i-j). These seeds on germination showed a variable degree of viability. The viable seeds albeit in reduced number were propagated to the next generation.

**Figure 3.**
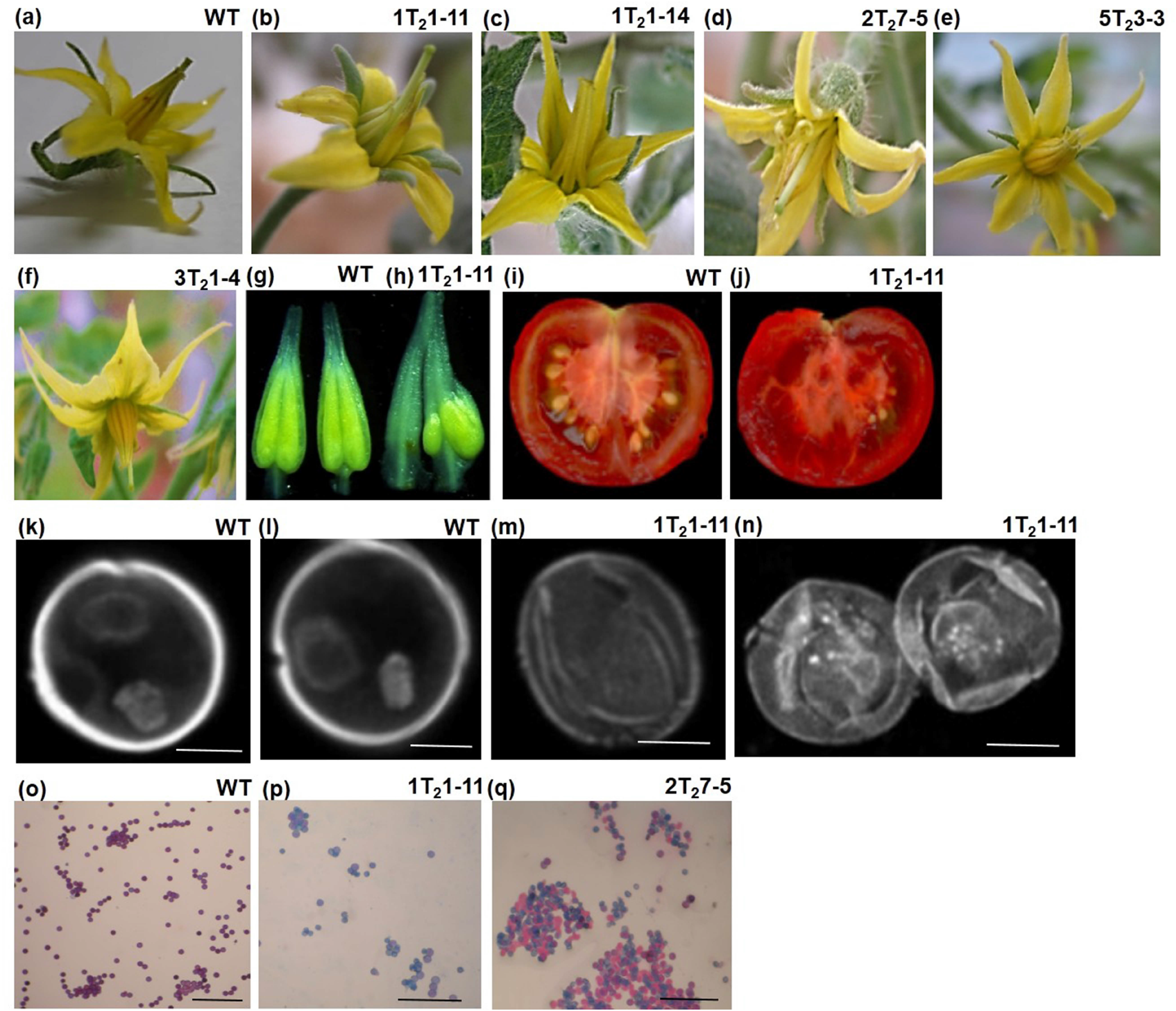
Abnormal flower, anther, fruit and pollen morphology manifested by *MSH2*-RNAi T_2_ Plants. (a) WT flower (b-d) Abnormal floral morphology, particularly in stamens, in strongly *MSH2* silenced lines. (e-f) Flowers of partially *MSH2* silenced lines, 5T_2_3-3 and 3T_2_1-4 are nearly similar to the WT. (g) Anther cones of line WT. (h) Abnormal anther cones of line 1T21-11. Note the absence of characteristic yellow coloration in 1T_2_1-11. (i-j) Vertical cut section of red ripe fruit showing highly reduced seed set in line 1T_2_1-11 (i) compared to WT (j). (k-l) WT pollen grains are bicellular with vegetative and generative cells compared to empty (m) and shriveled (n) pollen grain in line 1T_2_1-11 (K-N, Scale bar, 20 μm). (o-r) Alexander staining showing viable pollen grains in WT (o), line 1T_2_1-11 (p) 2T_2_7-5 (q) and 2T_2_5-5 (r). (l-o, Scale bar, 200 μm).

The examination of pollens revealed that strongly silenced lines produced pollens with the deformed shape. Unlike WT pollens that were round with a vegetative and generative cell (Figure 3k-l), the pollens of silenced lines showed the deformed shape and were either empty or contained the remnant of chromosome fragments (Figure 3m-n). It is likely that pollen grains collapsed either due to lack of nourishment in deformed anthers or *MSH2*-silencing affected some other gene.

The viability of WT and *MSH2*-RNAi pollens was examined by Alexander staining. In WT, all pollens exhibited a red-purple fluorescence, strongly reflecting the high male gametophytic fertility of these plants (Figure 3o). In contrast, *MSH2*-RNAi lines such as 1T_2_1-11 displayed mostly inviable pollens (Figure 3p-r). As a consequence, *MSH2*-RNAi plants set fruits bearing ca. 5% seeds. The tomato line 1T_2_1-11 with the highest reduction in *MSH2* expression showed a maximum loss in the fertility with > 90% blue stained pollen grains (Figure 3p). The fruits of 1T_2_1-11 line bore very few viable seeds along with underdeveloped seeds (Figure 3j).

### *MSH2*-RNAi lines display tetraploid meiosis progression

The deformities in the anther morphology are reported to be associated with defects in male meiosis in tomato (Rick, 1948; Jeong *et al*., 2014) and Arabidopsis (Panoli *et al*., 2006). Considering that observed stamen abnormalities could be associated with meiosis, we examined meiosis progression in the *MSH2* silenced lines. The pollen mother cells (PMCs) were isolated at different stages of floral bud development as outlined by Brukhin *et al*. (2003) from *MSH2* silenced lines and WT. A complete meiotic progression was examined for WT which progressed normally with characteristic landmarks for different stages of the meiosis, such as zygotene (Figure 4a), pachytene (Figure 4b), diplotene and diakinesis (Figure 4c-d) followed by metaphase (Figure 4e), early and late anaphase (Figure 4f-g), early and late telophase (Figure 4h-i).

**Figure 4.**
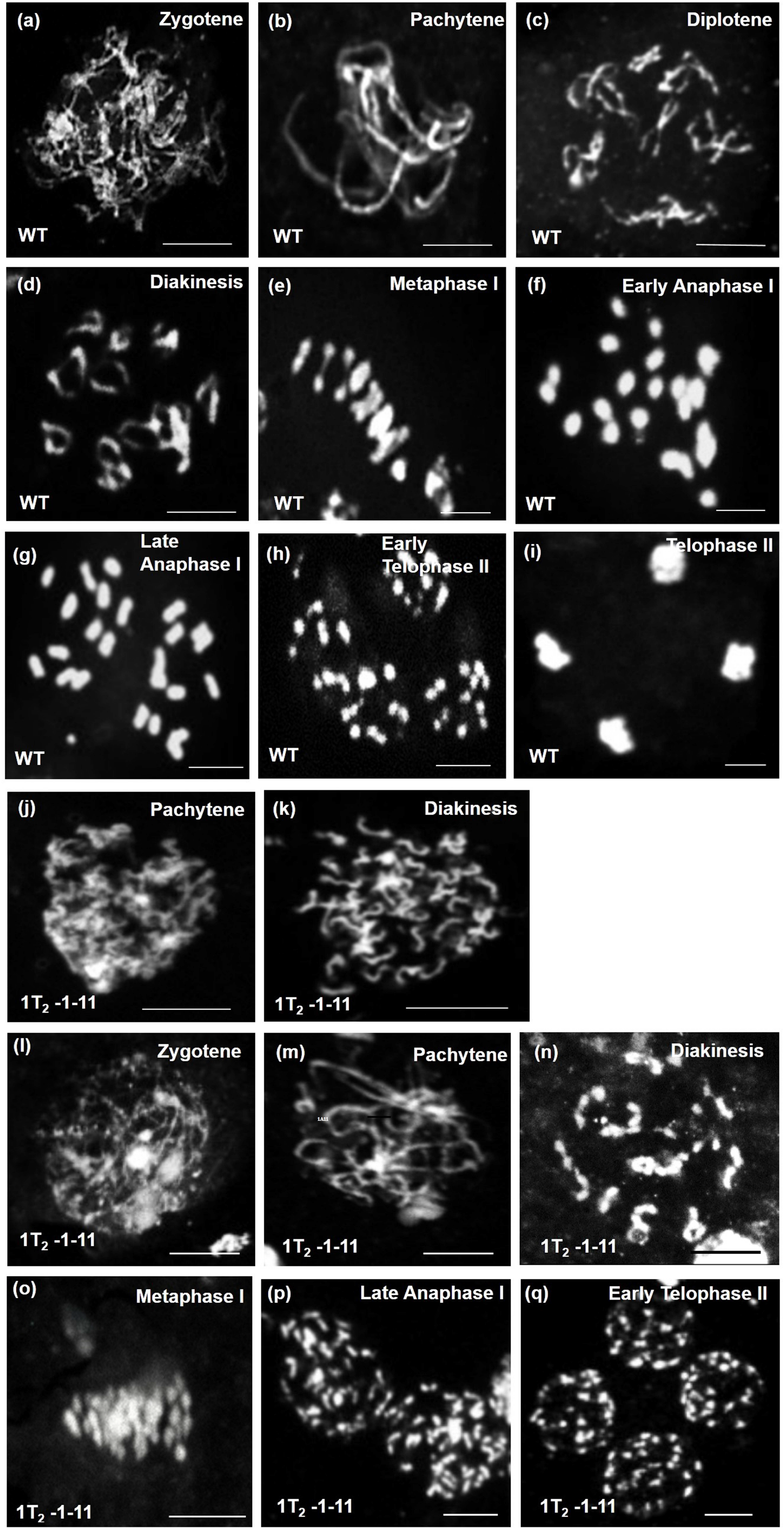
Male meiosis in wild-type and *MSH2*-RNAi tomato lines. Representative meiotic stages of wild-type (a–i) and *MSH2*-RNAi (j-q) lines from zygotene to telophase. In *MSH2*-RNAi meiocytes, few univalent chromosomes were observed that were assumed to be in early prophase I, (j-k). A proportion of male meiocytes of *MSH2*-RNAi lines underwent tetraploid meiosis that progressed from prophase to telophase (l-q). Note the formation of diploid tetrads (q). Scale bar, 10 μm.

In strongly silenced *MSH2*-RNAi lines, irrespective of the stage of floral bud development, predominant short condensed chromosomes were observed in the majority of the meiocytes. These meiocytes were either arrested at the prophase I (Figure 4j-k) or completed later stages of meiosis (Figure 4l-q). The abnormal increase in the number of chromosomes in meiocytes was suggestive of either premature loss of sister chromatid cohesion or increase in ploidy leading to the formation of diploid tetrads ((Figure 4q). In these lines, very few meiocytes were observed with WT complement of chromosomes.

To quantify the chromosome number in meiotic cells of *MSH2*-RNAi lines, we used Centromere Protein C (CENP-C) antibody that specifically labels the centromere. Immunolabelling of WT prophase I meiocytes with CENP-C antibody showed 12 centromeres at pachytene indicating the likely presence of 12 bivalent chromosomes (Figure 5a-c). In contrast, the meiocytes of *MSH2* silenced line at a stage equivalent to prophase I showed 48 centromeres (Figure 5d-f). There are two possibilities to explain the presence of 48 centromeres. First, a loss of sister chromatid cohesion in the prophase I may lead to the appearance of 48 sister chromatids. Second, an increase in ploidy to tetraploid in the pollen mother cell followed by an absence or delay in sister chromatid cohesion may result in 48 chromosomes. Analysis of meiotic stages favors the second scenario where we can observe the occurrence of diploid tetrads where the sister chromatids separate during anaphase II. Consistent with above, among the meiocytes examined, most contained 48 chromosomes as signified by CENP-C, however, in few meiocytes, the meiosis proceeded to anaphase I with segregation of 24 chromosomes to each pole (Figure 5g-i). Beyond this, other stages of meiosis I and meiosis II was rarely observed indicating a block in the meiosis of the silenced lines. However, a few cells displayed 12 centromeres indicating that silenced lines generate haploid spores albeit at highly reduced level.

**Figure 5.**
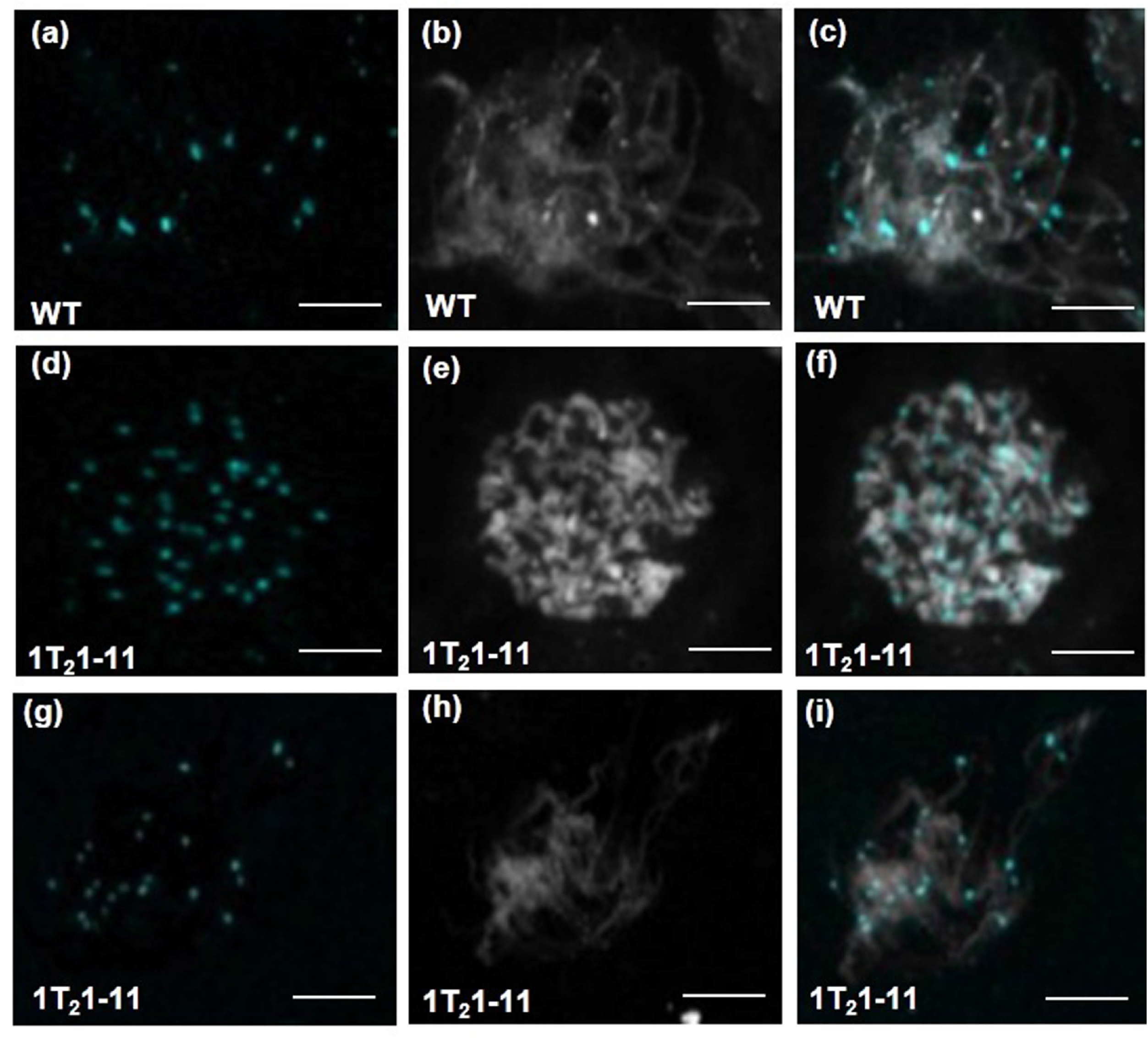
Immunolocalization of centromere marker protein-CENPC in nuclei of WT and *MSH2*-RNAi, 1T_2_1-11 line during prophase I. For each nuclei, the CENPC and DAPI staining along with merged signals is shown. (a-c) Zygotene nuclei with 12 centromere spots are likely indicating 12 bivalents in WT (a) with DAPI (b) and merged image (c). (d-f) Assumed zygotene nuclei with 48 centromere spots in 1T_2_1-11 line (d) are likely indicating 48 univalents along with DAPI (e) and merged image (f). (g-i) Assumed zygotene nuclei with 24 centromere spots in line1T_2_1-11 (g) likely indicating the presence of 24 bivalents along with DAPI (h) and merged images (i). The nuclei were examined using a 63X (1.4 N.A.) oil-immersion objective on a Leica SP5 confocal microscope. Scale Bar, 10 μm.

We observed that only a few tetraploid cells completed meiosis II with the formation of diploid tetrads at the end of meiosis II (Figure 4k-o). The analysis of relative proportions of meiocytes revealed that in 1T_2_1-11 line, 25% meiocytes displayed abnormal tetraploid chromosomes, 67% tetraploid meiocytes and 8% displayed diploid meiosis (Table 2). Similarly, the 2T_2_7-5 line had 25% meiocytes with abnormal tetraploid and 75% tetraploid nuclei giving rise to diploid microspores. Consistent with this, the anthers of *MSH2* silenced lines contained largely empty shrunken pollen at the end of meiosis. Taken together these results confirm that *MSH2*-RNAi meiocytes display abnormal meiotic progression resulting in tetraploid meiocytes. The tetraploid meiocytes were observed only in 1T_2_1-11 and 2T_2_7-5 lines with high silencing of *MSH2* transcript. The 3T_2_1-4 line with moderate *MSH2* silencing and 5T_2_3-3 line with no *MSH2* silencing displayed diploid meiocytes similar to WT (Figure S6).

**Table 2:** Number of male meiocytes in different *MSH2*-RNAi lines with reference to WT and the number of seeds per fruit in the respective lines.

| **Line No** | **Total No. of meiocytes count** | **No. of Diploid meiocytes** | **No. of Normal Tetraploid meiocytes** | **No. of Abnormal Tetraploid meiocytes** | **No. of seeds per fruit** |
| --- | --- | --- | --- | --- | --- |
| WT | 60 | 60 | 0 | 0 | 40-50 |
| 1T<sub>2</sub>1-11 | 60 | 5 | 40 | 15 | 0-5 |
| 2T<sub>2</sub>7-5 | 60 | 0 | 45 | 15 | 2-10 |
| 3T<sub>2</sub>1-4 | 60 | 60 | 0 | 0 | 40-50 |
| 5T<sub>2</sub>3-3 | 60 | 60 | 0 | 0 | 40-50 |

### Analysis of T-DNA integration sites in *MSH2*-RNAi lines using FPNI-PCR

We also examined whether the tetraploid meiocytes or defect in cytokinesis was caused by inactivation of some important gene due to insertion of the transgene. To locate the site of transgene insertion, FPNI-PCR was performed on genomic DNA of 1T_2_1-11 and 2T_2_7-5 tetraploid *MSH2*-RNAi lines. The amplified PCR product flanking the right border of inserted T-DNA was cloned and sequenced. The flanking sequence of line 1T_2_1-11 showed 98% homology to *Tic22* gene in chromosome No. 9 (Table S1, Figure S7a,b) and that of 2T_2_7-5 showed hit in *TRM32* gene, a member of the TON1 Recruiting Motif superfamily (Table S1, Figure S7c,d). While TRM1 reportedly associates with cortical microtubules, not all TRM proteins associate with microtubules. Moreover, the sequence diversity among the TRM family members with varied transcriptional expression patterns indicates that different members may have different roles (Struk and Dhonukshe, 2014). Considering that both 1T_2_1-11 and 2T_2_7-5 *MSH2*-RNAi lines show tetraploidy, it is unlikely that insertional inactivation of these genes leads to tetraploidy in the silenced lines.

### Root tip cells show diploid nuclei with sporadic defect in cell plate formation

In the present study, *MSH2*-RNAi tomato lines generated tetraploid meiocytes. However, no predominant enlarged organ size, a characteristic of increased ploidy was observed during the vegetative growth of plants. Even though the silenced lines displayed abnormal floral organs, these too showed no significant enlargement of floral structures. To ascertain whether *MSH2* silencing leads to tetraploidy in the progeny of the plants, we examined somatic chromosomal count in root tips of silenced and control seedlings. The nuclei in WT root tips displayed diploid chromosome number. Interestingly the *MSH2*-RNAi lines 1T_2_1-11 and 2T_2_7-5 with tetraploid meiocytes also displayed diploid chromosomal nuclei in root tip tissues indicating that these plants were diploid. At the same time, few cells showed no intervening cell wall between the nuclei suggesting that *MSH2* silencing affected the cell plate formation (Figure 6). The loss of intervening cell wall resulted in the formation of several syncytial nuclei within a cell (Figure 6e,f). In contrast, the weakly silenced *MSH2*-RNAi lines with diploid meiocytes exhibited root tip with uniformly sized diploid nuclei with distinct cell plate formation (Figure 6c) indicating that loss of cell plate was a feature specific to lines with a higher degree of *MSH2* silencing.

**Figure 6.**
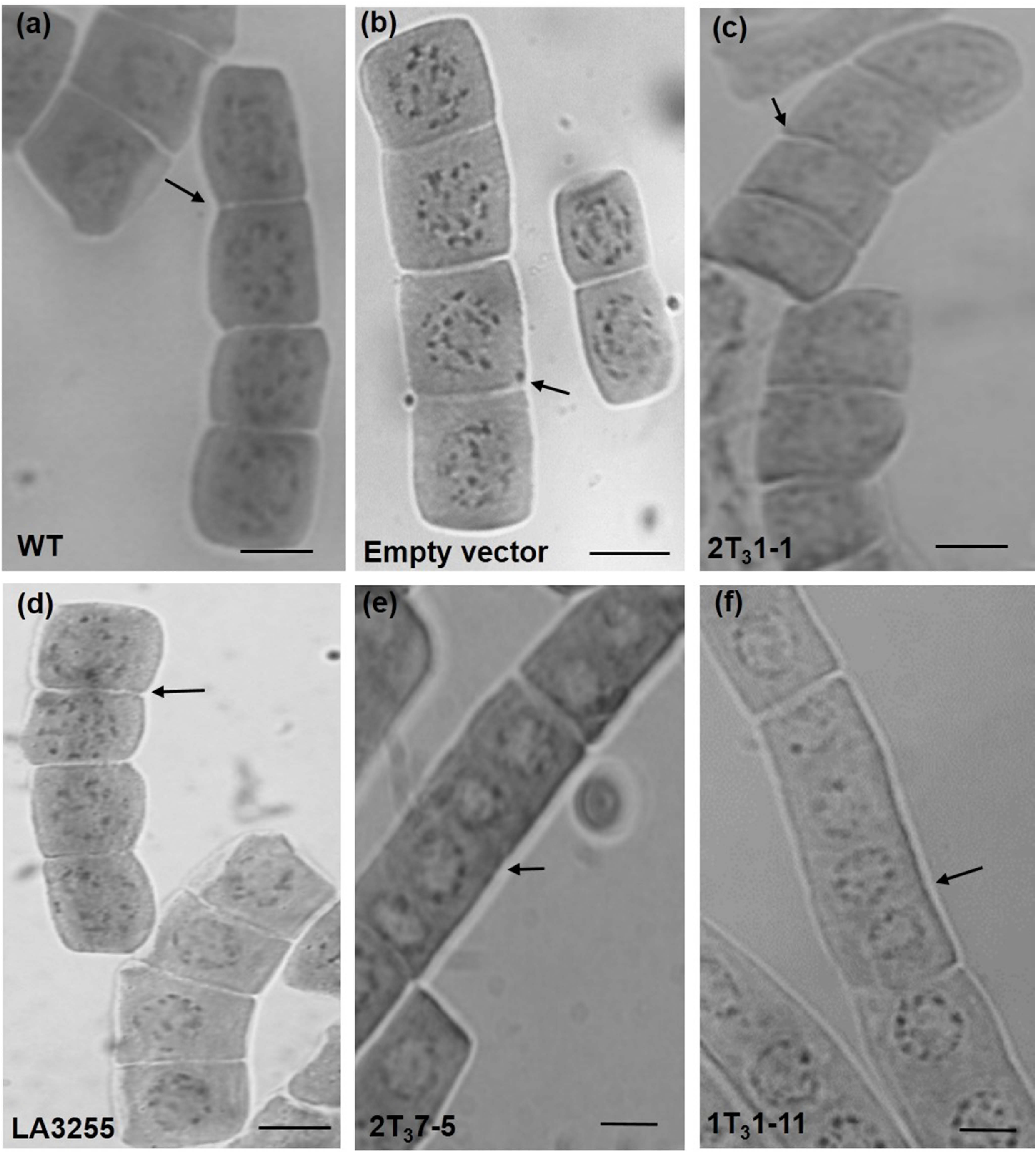
Mitosis in WT and T_3_ *MSH2*-RNAi lines root tip cells of progenies of (a-d). WT (a), empty vector control lines (b), Progenies of *MSH2*-RNAi, 2T_2_1-1 line bearing diploid meiocytes (c), and tetraploid line LA3255 (d) showing clear separation of the diploid nucleus in mitotic cells of root tip by intervening cell plate. (e-f). Progenies of strongly silenced *MSH2*-RNAi line 2T_2_5-1 (e) and 1T_2_1-11 (f) with tetraploid meiocytes show the absence of cell plate between dividing mitotic nuclei. Scale Bar, 100 μm.

To confirm whether loss of cell plate formation was related to the tetraploidy, we examined root tips of tomato tetraploid line, LA3255. The root cells of the LA3255 line showed normal tetraploid nuclei with no defect in the cell plate formation (Figure 6d). Consistent with tetraploid nature of the LA3255 line, the staining of root tip nuclei with DAPI showed larger nuclei in LA3255 line compared to diploid wild-type and MSH2-RNAi lines nuclei (Figure S8).

## DISCUSSION

The DNA mismatch repair is the highly conserved biological pathway that maintains the fidelity of DNA replication and genetic stability. In the mammalian system, the defects in MMR lead to abnormalities in meiosis and increases predisposition to certain types of cancer (Li, 2008). Consistent with the vital role of MMR, the *MSH2* in tomato appeared recalcitrant to mutagenesis, as evident by the absence of nonsynonymous mutations in M2 lines screened by TILLING. In the mammalian system, the polymorphism in *MSH2* significantly increases the risk of breast cancer (Hsieh *et al*., 2016). The paucity of polymorphism in *MSH2* as revealed by EcoTILLING of tomato accessions is in conformity with the indispensable nature of the gene. Though a missense and few silent mutations were identified, these had no influence on the protein function as predicted by SIFT.

In plants, the role of *MSH2* gene in MMR has been mainly inferred based on its homology with yeast and mammalian genes. In Arabidopsis, a *MSH2* T-DNA insertional mutation accumulates mutations in microsatellites and other loci indicating a role of *MSH2* in MMR (Leonard *et al*., 2003; Hoffman *et al*., 2004). MSH2 reportedly plays a role in repair of the UV-B induced DNA damage in the mammalian system, (Peters *et al*., 2003; Seifert *et al*., 2008). In plants as well, MSH2 likely plays a role in the repair of UV-B induced Cyclobutane Pyrimidine Dimer (CPD). The Arabidopsis *MSH2* mutant on UV-B exposure shows enhanced formation of CPD suggesting a role of MSH2 in the repair of UV-B damage (Lario *et al*., 2011). Similarly, in strongly silenced tomato *MSH2*-RNAi lines, thymine dimer formation is exacerbated by exposure to UV-B. Also, *MSH2*-deficient plants show inefficient repair of thymine dimers even on exposure to visible light that stimulates photoreactivation repair. From the preceding, it is apparent that strongly silenced *MSH2*-RNAi lines have low *in vivo* MMR activity as evident by enhanced UV-B damage.

The propagation of Arabidopsis *MSH2* T-DNA insertional mutant line revealed the accumulation of a wide variety of mutations such as morphological abnormalities, reduction in seed/silique development and loss of germination efficiency (Hoffman *et al*., 2004). Analogously, strongly silenced tomato *MSH2*-RNAi lines also show multiple abnormalities throughout the plant development, mainly during floral development with abnormal anthers. The lines with abnormal stamens showed reduced fruit set and highly reduced seed numbers in the fruits. In contrast to Arabidopsis *MSH2* mutant lines where seed set on average was reduced to only 50% (Hoffman *et al*., 2004), in tomato silenced lines, it was less than 10%.

The reduction in seed set can result from the failure in the embryo development. Considering that the *MSH2* silenced lines developed abnormal anthers, reduced seed set may have resulted from the failure to make viable pollens in the deformed anthers. Consistent with this, Alexander staining showed nonviable pollens with blue staining in highly silenced lines. Since WT and moderately silenced lines showed viable pollens with red staining, it can be assumed that *MSH2* silencing leads to loss of pollen viability in tomato. Thus the reduced seed set in the silenced lines is in conformity with the loss of pollen viability.

The successful pollen development in plants requires operation of proper meiosis in the developing anther (Ma, 2005). In several plants defects in the anther formation are associated with defective meiosis or meiotic arrest (Walbot and Egger, 2016). In tomato, heat-induced abnormal anther development in a conditional male sterile mutant is associated with reduced pollen set and loss of pollen viability (Rick and Boynton, 1967). It is, therefore, likely that silencing of MSH2 expression affected anther development that in turn may have affected the progression of meiosis.

Consistent with this view, the *MSH2* silenced lines showed a meiotic arrest in the anthers. The majority of pollen mother cells (PMC) of *MSH2*-RNAi lines fail to complete the meiotic cycle. These PMCs initiate meiosis as evident by the onset of zygotene. However, the meiosis gets stalled at the final stage of prophase I – diakinesis, as the chromosomes are at their most condensed form. The stalling at diakinesis is seemingly related to the increase in the ploidy as CENP-C stained cells showed the presence of 48 univalent chromosomes. The stalling of the meiosis is not total, as few CENP-C stained cells also showed 24 bivalent chromosomes, which completed meiosis presumably generating diploid pollens. However, most pollens were shrunken, and only a few showed Alexander staining. It is plausible that loss of pollen viability was largely for the diploid pollens. From the preceding, it is likely that incomplete meiosis and the formation of nonviable pollens contribute to the reduced seed set in the strongly-silenced *MSH2*-RNAi lines.

The process that generates tetraploid nuclei in the PMC remains to be investigated. In several plants including tomato during microsporogenesis, the nuclei can transfer to the neighboring cell by cytomixis (Weiling, 1965; Mursalimov *et al*., 2013). Such intercellular transfer of nuclei can influence the ploidy level of the produced pollens. However, such cytomictic transfer of nuclei most probably does not contribute to tetraploid nuclei, as in tomato cytomixis is observed only ca. 15% of PMC (Weiling, 1965). The occurrence of a large number of polyploid pollen mother cells does not favor cytomixis to be the cause of tetraploid nuclei. The probability that tetraploid meiocytes arise due to *MSH2*-silencing-induced endomitosis also seems remote. In Chrysanthemum, the fusion of two PMC forms tetraploids, albeit this process occurs at very low frequency (Kim *et al*., 2009), and cannot account for high numbers of tetraploid meiocytes in *MSH2* silenced lines.

The root cells of *MSH2*-RNAi-silenced lines show defective cytokinesis with multiple nuclei in syncytial cells. The similar syncytial cells may also form due to defective cytokinesis during meiocytes archesporial cells development. It is likely that the fusion of nuclei in syncytial cells gives rise to the tetraploid nuclei in PMC of *MSH2*-RNAi-silenced lines. The fusion of nuclei resulting in the stochastic formation of tetraploid PMC initials has been documented in mutants defective in cell plate formation. In Arabidopsis, the mutations in sterol methyl transferase and callose synthase lead to defect in cell plate formation predominantly in the floral tissues. Both mutants also form tetraploid meiocytes and diploid gametes in both male and female sporogenesis (de Storme *et al*., 2013). The formation of tetraploid meiocytes in *MSH2* silenced lines is analogous to tomato *pmcd1* mutant, where cytokinesis failure in meiocytes archesporial cells generates tetraploid nuclei (de Storme and Geelan, 2013a).

The *MSH2* silenced lines show the formation of the very few viable pollens and the progeny of these plants were exclusively diploid. The strongly reduced seed set in silenced lines seems to indicate that only diploid embryos survived post-fertilization. Another reason could be defective endosperm development due to unbalanced 1: 1 ratio rather than normal 1:2 ratio of the paternal and maternal genome (Scott *et al*., 1998). The absence of triploid progeny may also result from the existence of strong triploid block in members of Solanaceae including tomato (Ehlenfeldt and Ortiz 1995; Nilsson, 1950). The absence of triploid or tetraploid progeny in *MSH2*-RNAi lines is also consistent with the reported absence of triploid/tetraploid plants in progeny of tomato *pmcd1* mutant that shows a cytokinesis defect and generates diploid pollens (de Storme and Geelan, 2013a)

The pre-meiotic doubling of chromosome occurs in unisexual and parthenogenetically reproducing animal species (Itono *et al*., 2006). It is believed that in higher plants such chromosomal doubling rarely occurs (de Storme and Geelan, 2013b). In higher plants, the doubling of chromosomes reportedly occurs after the onset of meiosis via meiotic restitution (de Storme and Geelan, 2013b). Considering that doubling of chromosomes in *MSH2* silenced line occurs before the onset of meiosis, it is of a novel nature. The very high frequency of tetraploid meiocytes also hints for a likely function of *MSH2* in the regulation of chromosomal doubling via a mechanism that remains to be deciphered. Since the formation of tetraploid meiocytes in *MSH2* silenced line is quite similar to *pmcd1* mutant, the failure of cytokinesis appears to be the main contributor to this phenomenon.

In plants, MSH2 heterodimerizes with MSH3, MSH6, and MSH7. However, none of these heterodimers reportedly have any function in the regulation of meiosis. Conversely, the *MSH4* and *MSH5* mutants in Arabidopsis and rice though exhibit normal vegetative growth, show severe fertility reduction likely related to meiotic defects (Higgins *et al*., 2004; Lu *et al*., 2008; Luo *et al*., 2013; Wang *et al*., 2016). The MSH2-silenced lines also affected *MSH5* and *MSH7* transcript levels which may have affected the process of meiosis. The manifested tetraploid meiocytes formation and later failure in meiosis might be due to the reduction of these three transcripts. Alternately, the increased microsatellite instability caused due to low activity of the MSH2 protein may contribute to defective cytokinesis and tetraploid meiocytes formation.

Similar to *MSH2* silencing, loss of function lines in 40S ribosomal protein S6 kinases in Arabidopsis generate tetraploid PMCs and show a high degree of pollen abortion (Henriques *et al*., 2010). Likewise, mutations in glucan synthase-like 8 and sterol methyl transferase 2 lead to defects in somatic cell plate formation and the occasional fusion of syncytial nuclei (Chen *et al*., 2009; Guseman *et al*., 2010). It is, therefore, likely that T-DNA insertion in a meiosis/cytokinesis related gene may have caused the tetraploidy. The examination of two silenced lines indicated insertion in *TIC22* and *TRM32* genes; both of these genes have no reported role in meiosis or cell plate formation. Despite having an insertion in widely unrelated genes, both the silenced lines showed identical phenotype suggesting a close correlation between *MSH2* silencing and tetraploidy.

The tetraploid formation in *MSH2* silenced lines is more akin to the proposal that DNA replication or repair defects might produce signals that block cytokinesis and yield tetraploids in cancer cells (Ganem *et al*., 2007). While it is not reported in plants, in mice embryo fibroblasts, deficiency of *MSH2* in conjunction with *P53* revealed that MSH2 plays a role in preventing polyploidization of cells (Strathdee *et al*., 2001). In mammalian cells, *MSH2* has been identified as a target of the E2F transcription factors (Ren et al., 2002) wherein the loss of *MSH2* inhibits the interaction of *E2F* with *MSH2*, thus affecting the *E2F* signaling. It is believed that disruption of the *E2F* signaling affects the progression of cell cycle and promotes failure of cytokinesis. Since *MSH2* have a probable role in mammalian cytokinesis, it is plausible that *MSH2* may affect cytokinesis in plants in a similar fashion. Nonetheless, it remains to be determined whether *MSH2* alone or with its interacting partners plays a direct or an indirect role in regulation ploidy and cytokinesis in plants. It is also likely that *MSH2* silencing leads to wide ramification in plant cells and observed change in the ploidy level reflect its downstream effect.

In summary, our study indicates that *MSH2* silencing leads to tetraploid meiocytes formation in tomato. Additionally, *MSH2* silencing also affects cytokinesis with multiple nuclei in syncytial cells. Though the precise mechanism of *MSH2* action remains to be deciphered, these phenotypes indicate an additional role for *MSH* gene family. Our results also suggest that similar to cytomixis, the formation of syncytial cells may contribute to the tetraploid formation in higher plants. Uncovering the role of MSH2 in the regulation of cytokinesis and nuclear fusion will be of great interest in future studies and may contribute to an alternate route to the polyploid formation.

## EXPERIMENTAL PROCEDURES

### Screening for SNPs/mutations in *MSH2* gene

The population used for the EcoTILLING was same as described earlier in Mohan *et al*. (2016) and Upadhyaya *et al*. (2017). Genomic DNA was isolated from tomato accessions or EMS-mutagenized M_2_ lines following the procedure of Sreelakshmi *et al*. (2010). *MSH2* gene sequence was obtained from SGN (solgenomics.net, Solyc06g069230). CODDLE software (blocks.fhcrc.org/ proweb/ coddle) was used to predict the most deleterious region of the gene. The primers for SNP/mutation screening were designed using Primer3web version 4.0.0 (bioinfo.ut.ee/primer3). (Table S2). The SNP/mutation detection and haplotype assignment were carried out as described by Mohan *et al*. (2016). The sequence variations (Substitutions and Indels) were detected by using ‘Multalin’ software (http://multalin.toulouse.inra.fr/multalin/, Corpet (1988). The SNP frequency was calculated using the formula: (Total number of SNPs detected / total length of the screened fragment) X 1000 (Frerichmann *et al*., 2013). PARSESNP (Taylor and Greene, 2003) was used to analyze and display the variation in gene sequences, and SIFT (Sorting intolerant from tolerant; Sim *et al*., 2012) was used to predict the deleterious variations for protein functions.

### Cloning of a fragment of *MSH2* gene in pHELLSGATE 12 RNAi vector

For RNAi silencing, a 301 bp region in 4^th^ exon of *MSH2* showing no significant homology with any other tomato gene was selected. The 301 bp region from *MSH2* cDNA was PCR amplified using primers set with the attB sequence at the 5′ end (FP-AGATTCTCCAGTGATTGTTGCT; RP- CAGATCCTGTACCAAATCCCTA)̤ The PCR product was cloned into a pDONR201 vector with attP1 and attP2 sites using BP Clonase enzyme (*Invitrogen*). The gene fragment from the intermediate clone was transferred to the pHELLSGATE12 vector (CSIRO, Australia) by recombination between attL1/attL2 and attR1/attR2, mediated by LR Clonase enzyme (*Invitrogen*). The confirmed clone was transformed into *Agrobacterium tumefaciens* C58C1 strain.

### Generation of *MSH2*-RNAi lines

Tomato (cv. Arka Vikas) transformation was performed as described earlier (Sharma *et al*., 2009) yielding ten independent transgenic lines. The phenotyping of the transgenic plants was continually carried out in successive generations. The lines with 80-90% *MSH2* transcript silencing showed anther abnormalities. Therefore these lines were manually pollinated to propagate to next generation. Nonetheless, few T_2_ lines with very strong anther-defects did not survive as they produced insufficient viable pollens and failed to set fruits. The plants were carried forward till T_4_ generation. The following nomenclature was adopted for progressive generations; Line No. 2T_0_ indicates T_0_ line No.2, 2T_1_5 indicates T_1_ progeny No. 5 of Line No. 2, 2T_2_5-7 indicates its T_2_ progeny No. 7 of T_1_ plant No.5 of T_0_ Line No. 2.

### Southern blot analysis

Genomic DNA (~15 μg) of T_0_ transgenic lines was digested with BamHI and SalI to release a fragment size of 2.2Kb (Figure S3), while genomic DNA (~15 μg) of T_2_ transgenic lines was digested EcoRI or NotI or PacI (Figure S4) at 37°C overnight, separated on a 0.8% (w/v) agarose gel, and transferred to Hybond N^+^ membranes. αP^32^-dCTP labeled 2.2 Kb BamHI, and SalI digested (Figure S3) and *NPTII* (Figure S4) fragment was synthesized by PCR and used as the probe. The primers for amplification of *NPTII* probe amplify a 554 bp fragment in the coding region of *NPTII*. Prehybridization, hybridization, washing and developing of the blot was performed according to Sambrook and Russell (2001).

### RNA Isolation and Quantitative Real-Time PCR

RNA was isolated from young leaves with three independent replicates using the method as described earlier (Gupta *et al*., 2014). Complementary DNA (cDNA) was prepared from 2 μg of RNA using high-capacity cDNA Archive kit (Applied Biosystems, USA). Quantitative real-time PCR was also performed as described earlier by (Gupta *et al*., 2014). Gene-specific primers were designed from the sequences obtained from the SGN database using Primer 3 software (Table S3) while real-time data analysis were same as described by (Gupta *et al*., 2014).

### Analysis of in-vivo MSH2 activity

The *in-vivo* MSH2 activity was determined by measuring the amount of thymine dimer formation on exposure to UV-B light as described by Lario *et al*. (2011, 2015). Three samples of juvenile 6^th^ to 7^th^ node leaves were exposed for 6 h to UV-B radiation (2 Wm^−2^, TL 20 W/01-RS, Philips narrowband) in dark room at 22°C. Wild-type leaves were treated similarly albeit without UV-B radiation. One sample was frozen at the end of 6 h treatment. The second and third sample were transferred to the darkness and white light (100 μmol m^−2^ sec^−1^) respectively for 24 h. Samples were snap frozen and stored at −80°C till DNA extraction and thymine dimer determination.

The UV-B induced thymine dimers were quantified as described by Stapleton *et al*. (1993). Six μg of genomic DNA isolated from UV-B treated and control leaves was denatured, dot blotted onto Hybond N+ membrane (Perkin Elmer life Sciences, Inc.) and baked at 80°C. The membrane was blocked, washed and incubated with thymine dimer-specific monoclonal antibodies (Sigma-Aldrich, 1:2000 in TBS) for 3 h at room temperature with gentle agitation. The dimers were visualized by incubating with alkaline phosphatase conjugated secondary antibody (Sigma, 1:3000) and detection by nitroblue tetrazolium and 5-bromo-4-chloro-3-indolyl phosphate reaction. The bands were quantified by densitometry using Image J software. The thymine dimers were quantified using three independent biological replicates. However, in the figures, a single representative blot and the values derived from it using ImageJ software is shown.

### Meiotic Chromosome Preparation

Floral buds at the appropriate stages of the meiosis were picked according to Brukhin *et al*. (2003). The buds were fixed and stored in Carnoy’s solution [ethanol: chloroform: glacial acetic acid, (6:3:1)] at 4°C or −20°C. Fixed inflorescences were rinsed with Ethanol: glacial acetic acid (3:1) at defined intervals. Procedure for meiotic chromosome preparation was same as described by (Ross *et* al., 1997). Chromosomes were counterstained with AT-specific fluorescent dye, 4’-6-diamidino-2-phenylindole (DAPI, 1 μg/ml) (Sigma Laboratories). All images were captured using a 63X (1.4 N.A.) oil-immersion objective on a Leica SP5 confocal microscope. Images were further processed with Adobe Photoshop CS6.

### Mitotic Chromosome Preparation

Roots tips (1.5-2 cm) from 5-day-old seedlings were fixed in Carnoy’s solution for 24 h followed by washing with 70% (v/v) ethanol (15 min. 3X) and storage at 4°C. Procedure for mitotic chromosome preparation was same as described by (Babich *et al*., 1997). Then the tips were observed at 100X under Nikon compound microscope (Nikon Eclipse 80i) and photographed (Nikon NIS elements Basic version 3.1). Root tip nuclei were also assessed for nuclear size in somatic tissue by fixing in ethanol:acetic acid (3:1) fixative for at least 1 h, cleared in 70% ethanol and stained with 1μg/mL 4’,6-diamidino-2-phenylindole (DAPI) solution.

### Immunofluorescence

For immunofluorescence, synaptonemal complexes were prepared as described by Ross *et al*. (1997) and Chelysheva *et al*. (2010). 60% (v/v) acetic acid treatment was done at 45°C for 5 min with continuous stirring with a hook to remove maximum cytoplasm debris. Chromosomes were counterstained with DAPI and observed under the microscope to ensure proper slide preparation. The slides were microwaved in 10 mM citrate buffer pH 6 for 45s at 850 W for proper fixing of the chromosomes to slides. Slides were first immersed in 0.1M phosphate buffered saline (PBST): pH 7.4 containing 0.01% (v/v) TritonX-100 for 2×5 minutes. To block non-specific antibody binding, 100 μl of blocking buffer [1% (w/v) BSA in PBS] was applied directly to the slides. 100 μl of primary CENPC (Centromere protein C) antibody (1:100) in blocking buffer [PBS + 0.1% (v/v) Triton + 1% (w/v) BSA] was applied directly to the slides, covered with parafilm and incubated overnight at 4°C in a moist chamber. The slides were washed (2×5 minutes) with PBST before adding 100 μl of the secondary antibody. Secondary antibody, goat anti-rabbit 488 (Molecular Probes; diluted 1:500) was incubated for 90 minutes at room temperature. Finally, the slides were washed for 2×5 minutes and mounted in DAPI (10 μg/ml) in Vectashield antifade mounting medium. Slides were examined using a 63X (1.4 N.A.) oil-immersion objective on a Leica SP5 confocal microscope. Images were further processed with Adobe Photoshop CS6.

### Fusion primer and nested integrated PCR (FPNI-PCR)

To locate the unknown sequences flanking the insertion sites in the transgenic lines, FPNI-PCR described by Wang *et al*. (2011) with few modifications was used. The primers used for the FPNI-PCR are listed in Table S4. The cycling conditions were followed according to Wang *et al*. (2011). All PCR products were electrophoresed on a 2.0% (w/v) agarose gel and FPNI-PCR products showing sizes more than 500 bp were chosen for sequencing. The purified PCR product was sequenced using Sanger sequencing (Macrogen, South Korea). The sequences were aligned to the tomato genome sequence (Sol Genomics Network, https://solgenomics.net/) using BLAST program.

## ACKNOWLEDGEMENTS

The authors wish to thank Lorinda Anderson (Colorado State University, USA) for providing CENPC antibodies. This work was supported by International Atomic Energy Agency (IAEA) (grant no. 15632/R0-4 to RS), Department of Biotechnology, India (grant no. BT/PR/5275/AGR/16/465/2004; BT/PR/7002/PBD/16/1009/2012) to RS and YS, and University Grants Commission (research fellowship to SS).

## Authors Contributions

Conceived and designed the experiments, SS, AKP, RS, YS, MR. Performed the experiments, SS, AKP. Analyzed the data SS, AKP, RS, YS, and MR. Wrote the paper, SS, YS, RS and MR.

## SUPPORTING INFORMATION

Additional SUPPORTING INFORMATION may be found in the online version of this article.

**Figure S1.**
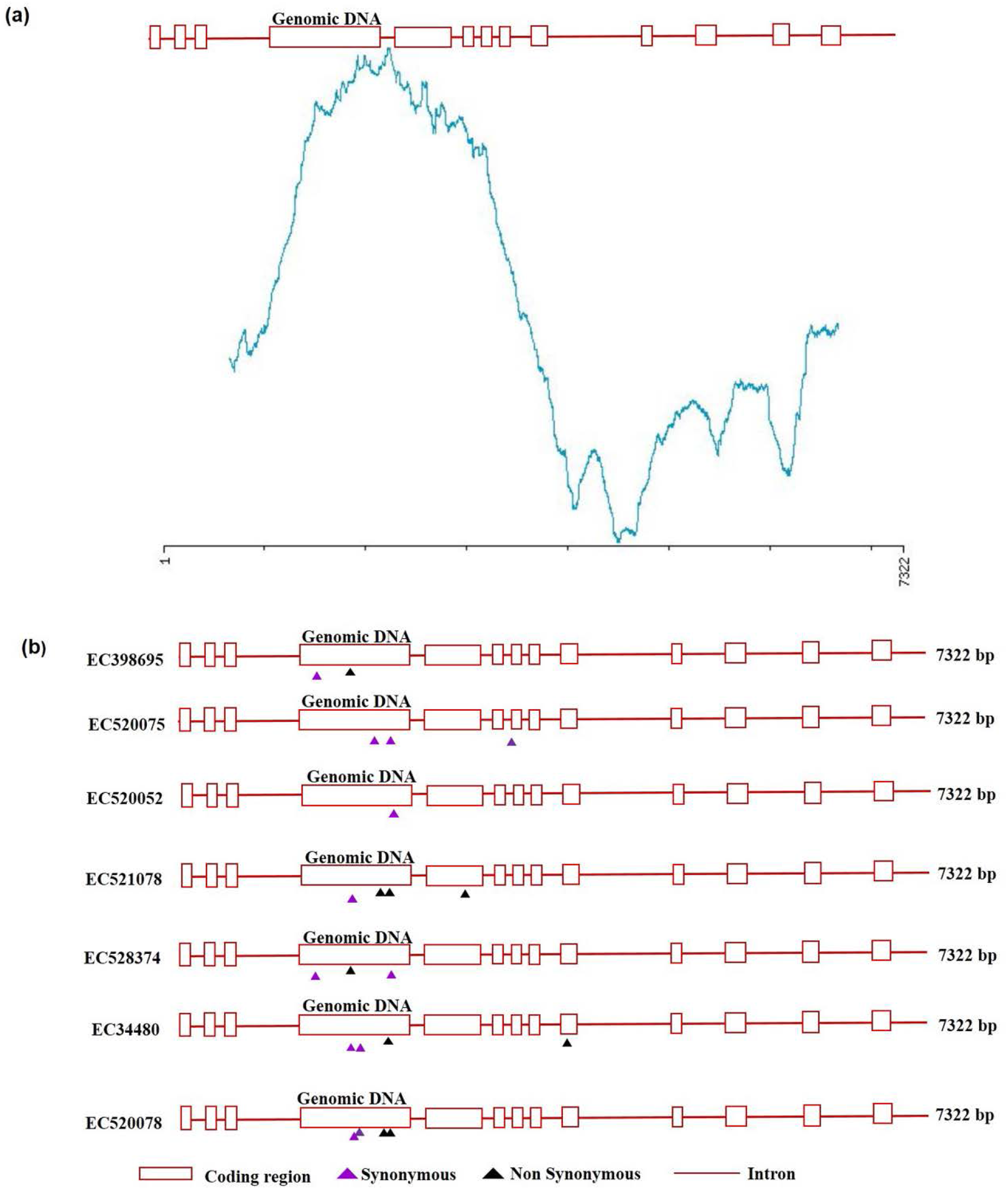
*In silico* prediction of most deleterious *MSH2* gene region by CODDLE (**a**) and distribution of SNPs detected in *MSH2* gene by EcoTILLING (**b**). (**a**) *In silico* prediction of most deleterious *MSH2* gene region by CODDLE. The tomato MSH2 gene consists of 7322 bp with 13 exons and 12 introns. The probability curve traced in blue represents the region of the gene where mutations would most likely deleteriously affect the function of encoded protein. Based on above prediction, the segment encompassing exons 3 to 9 was chosen for EcoTILLING. (**b**) Distribution of SNPs detected in *MSH2* gene by EcoTILLING. Red boxes denote exons interconnected with introns (solid red line). The black upright triangle indicates missense or nonsynonymous changes in the DNA sequence. The purple upright triangle indicates silent or synonymous changes. The numbers on the left and right of pictorial diagram respectively represent the accession number and total length of genomic DNA.

**Figure S2.**
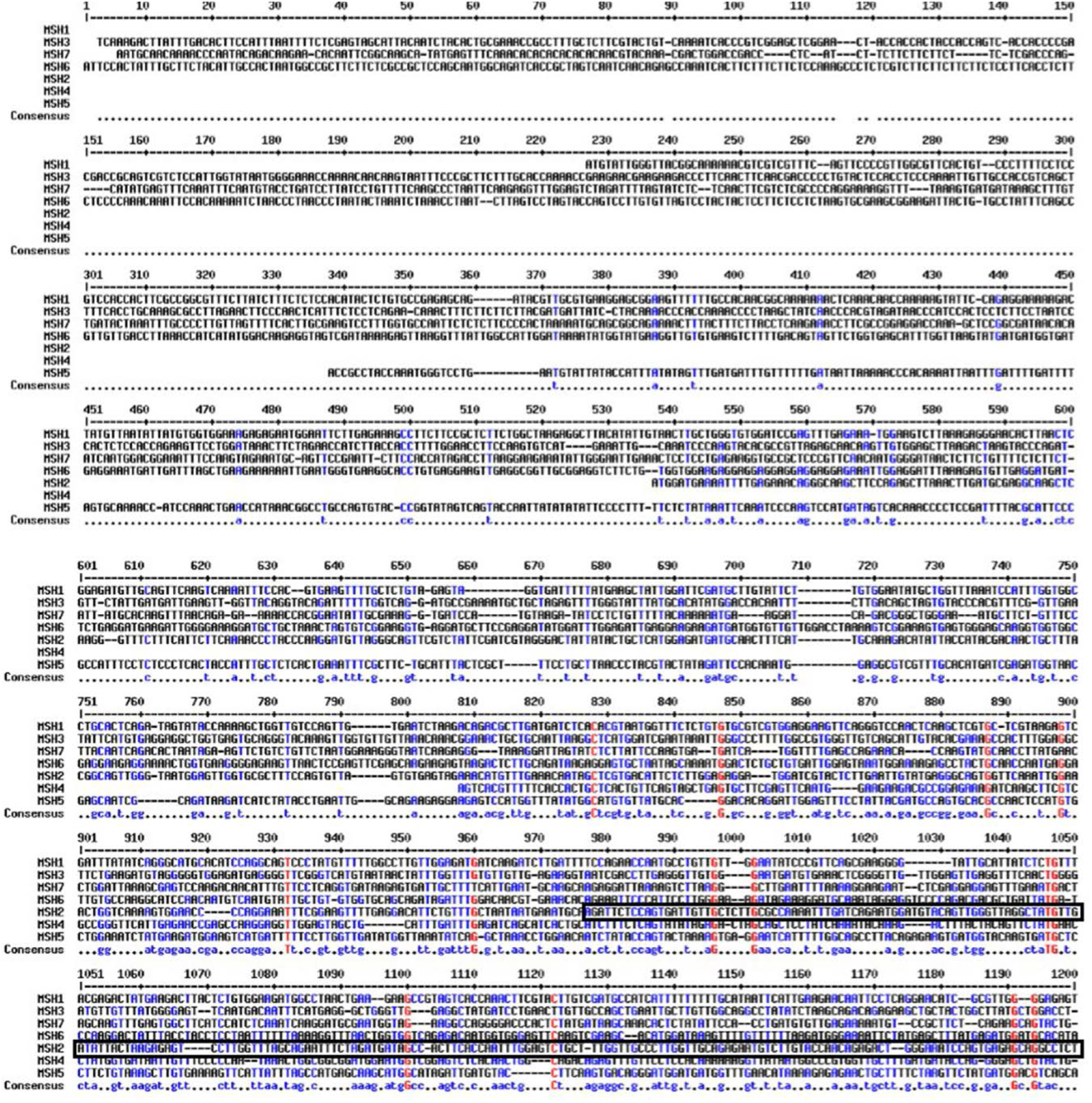
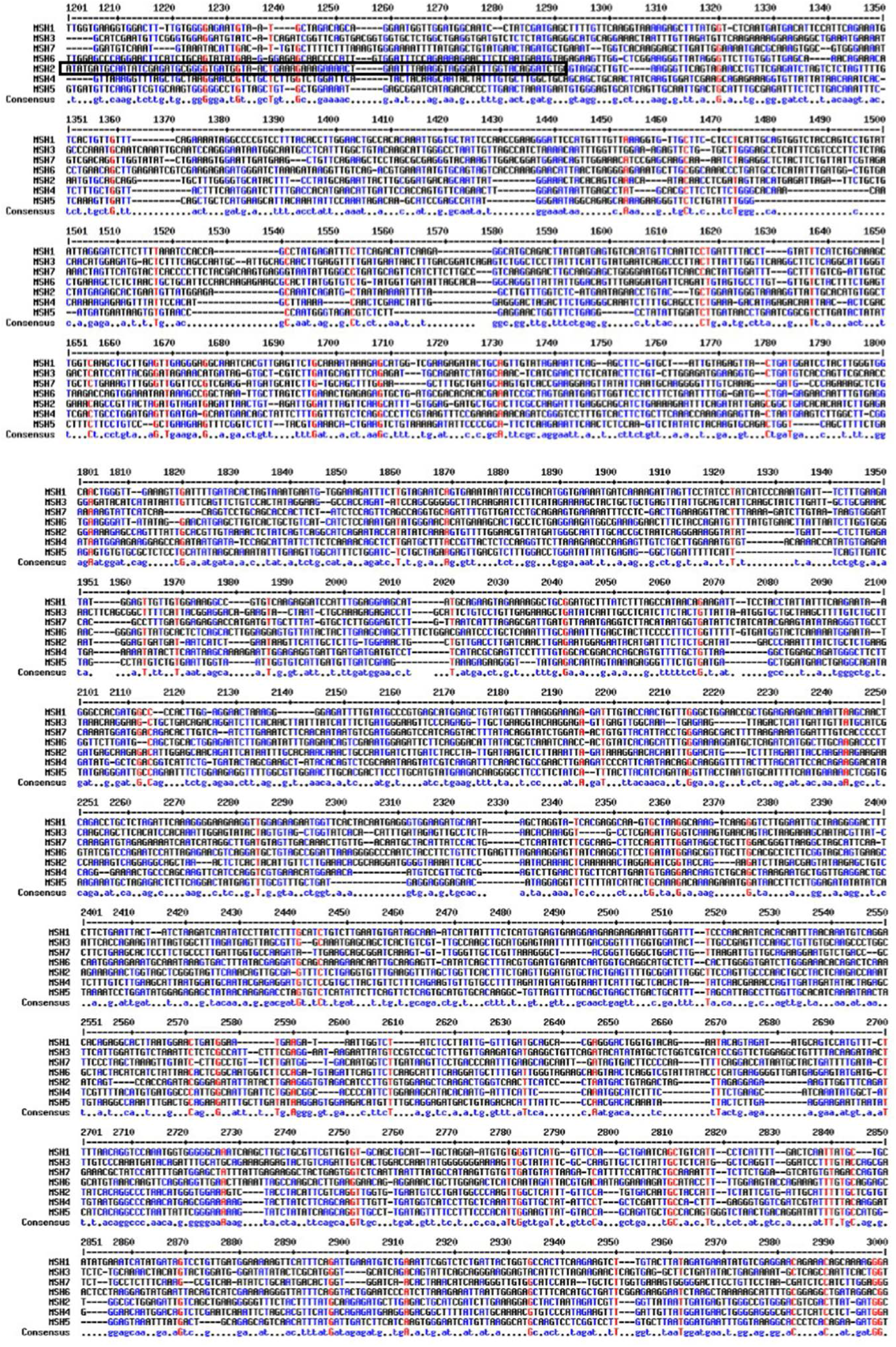
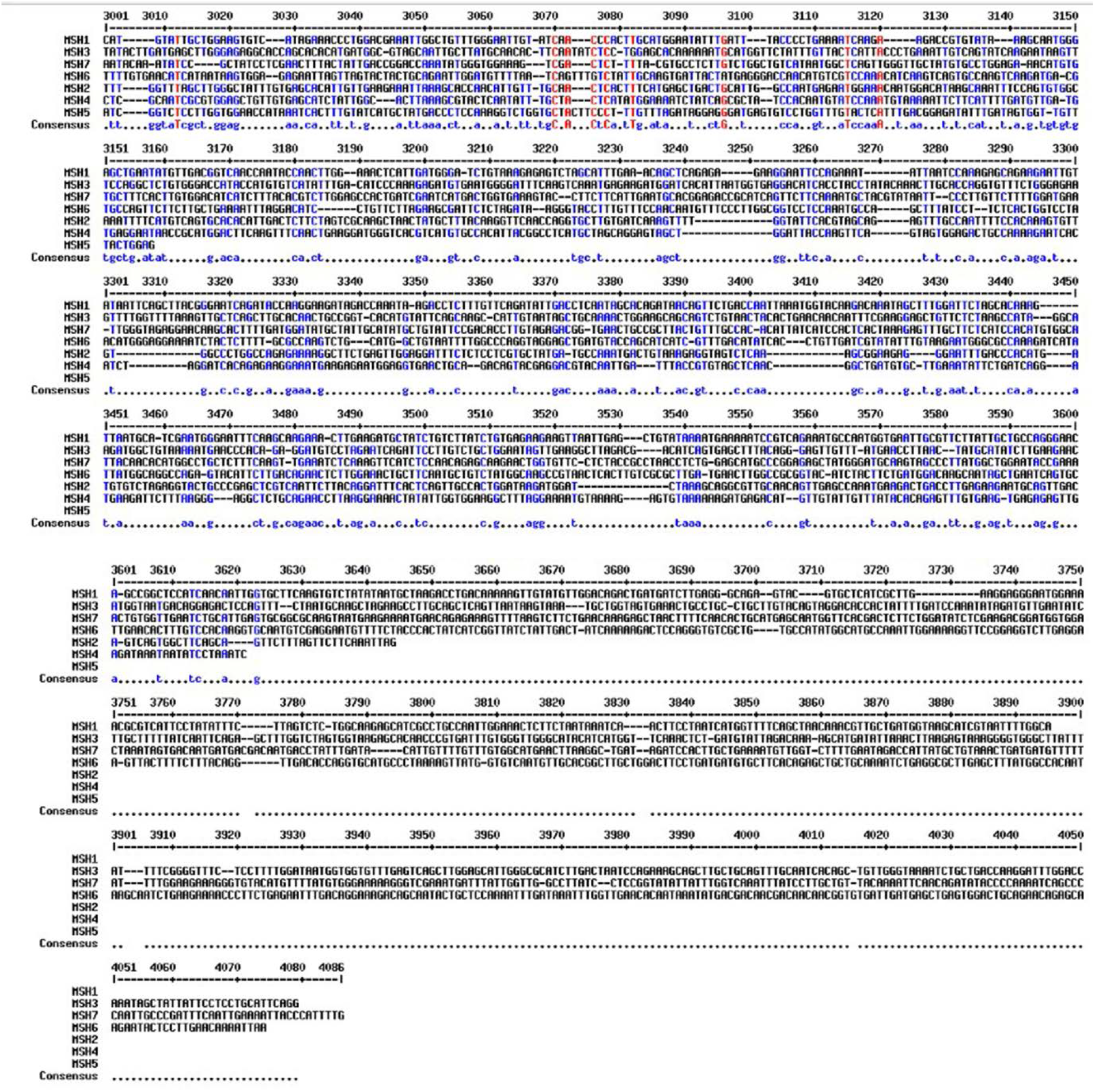
Multiple sequence alignment of mRNA sequences from tomato *MSH* family. The black box indicates the sequence of *MSH2* used for making *MSH2*-RNAi construct.

**Figure S3.**
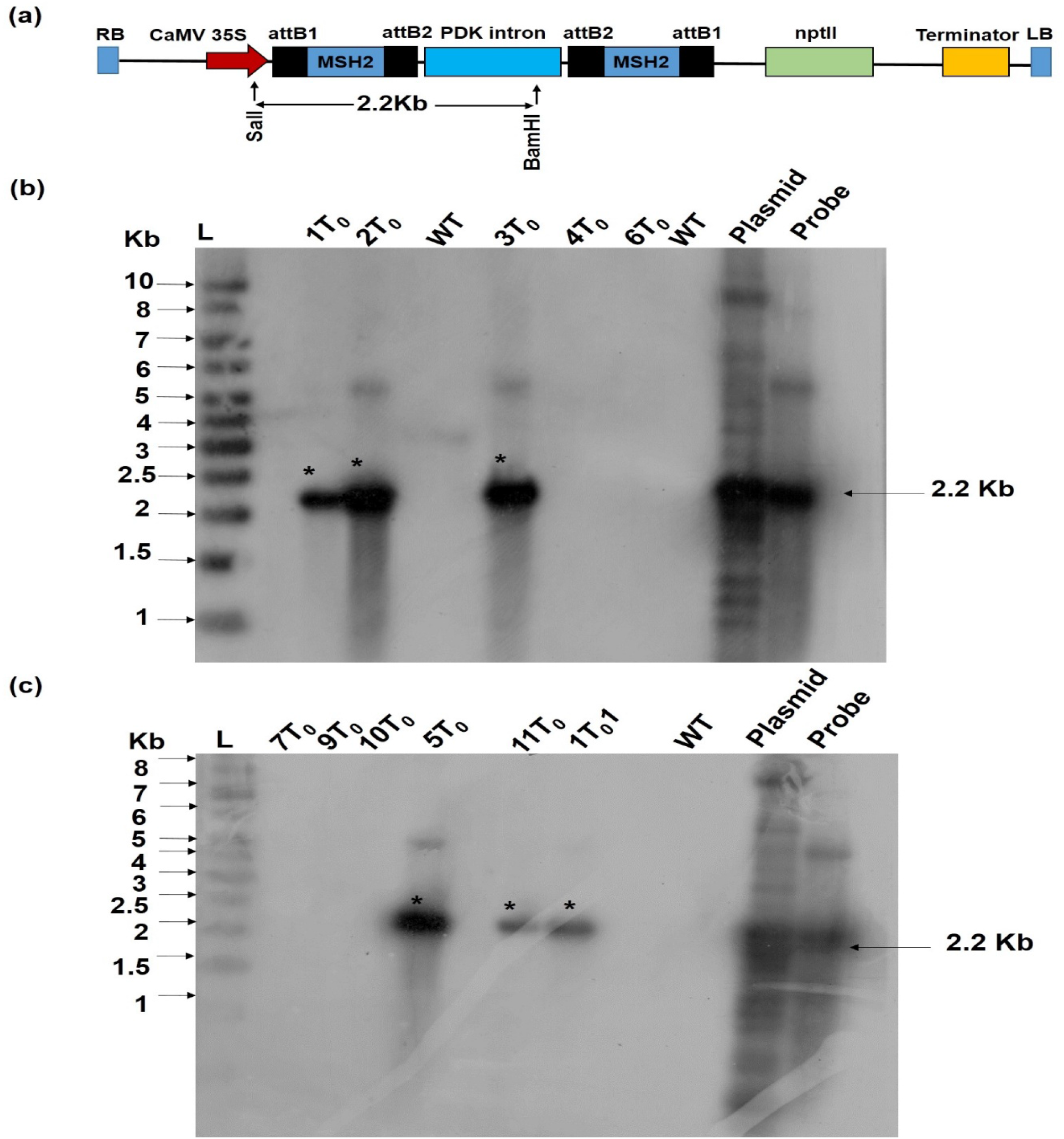
Generation of *MSH2*-RNAi transgenic tomato lines and Southern blot of T_0_ transgenic lines. (**a**) Schematic representation of the construct used for *MSH2* silencing. The construct contains one spliceable intron with the targeted *MSH2* sequence forming a hairpin when the construct undergoes transformation. *AttB1* and *attB2* represent the two short stretches of sequences that participate in the recombination reaction of the Gateway system. T35S indicates 35S terminator. The restriction sites for BamHI and SalI are also indicated. (**b-c**) Southern blot of T_0_ transgenic lines. Genomic DNA of T_0_ plants was digested with BamHI and SalI to release the insert. The Southern blot was probed with radiolabelled *NPTII–NOS* probe of 2.2 Kb size. Numbers on the top of lanes 1T_0_ to 6T_0_ in panel (**b**) and lanes 7T_0_ to 1T_0_ 1 in panel (**c**) indicate the plant number of respective T_0_ lines. **L**-1 Kb DNA Ladder (Fermentas). **WT**-Wild type genomic DNA (negative control); **Plasmid**-, Plasmid DNA bearing *MSH2*-RNAi construct. **Probe**-Radiolabelled NPTII–NOS 2.2 Kb probe. Note: The *NPTII–NOS* probe was obtained by digesting *MSH2*-RNAi plasmid with BamHI and SalI. The asterisk (*) indicates the presence of *NPTII–NOS* sequence in the transgenic lines.

**Figure S4.**
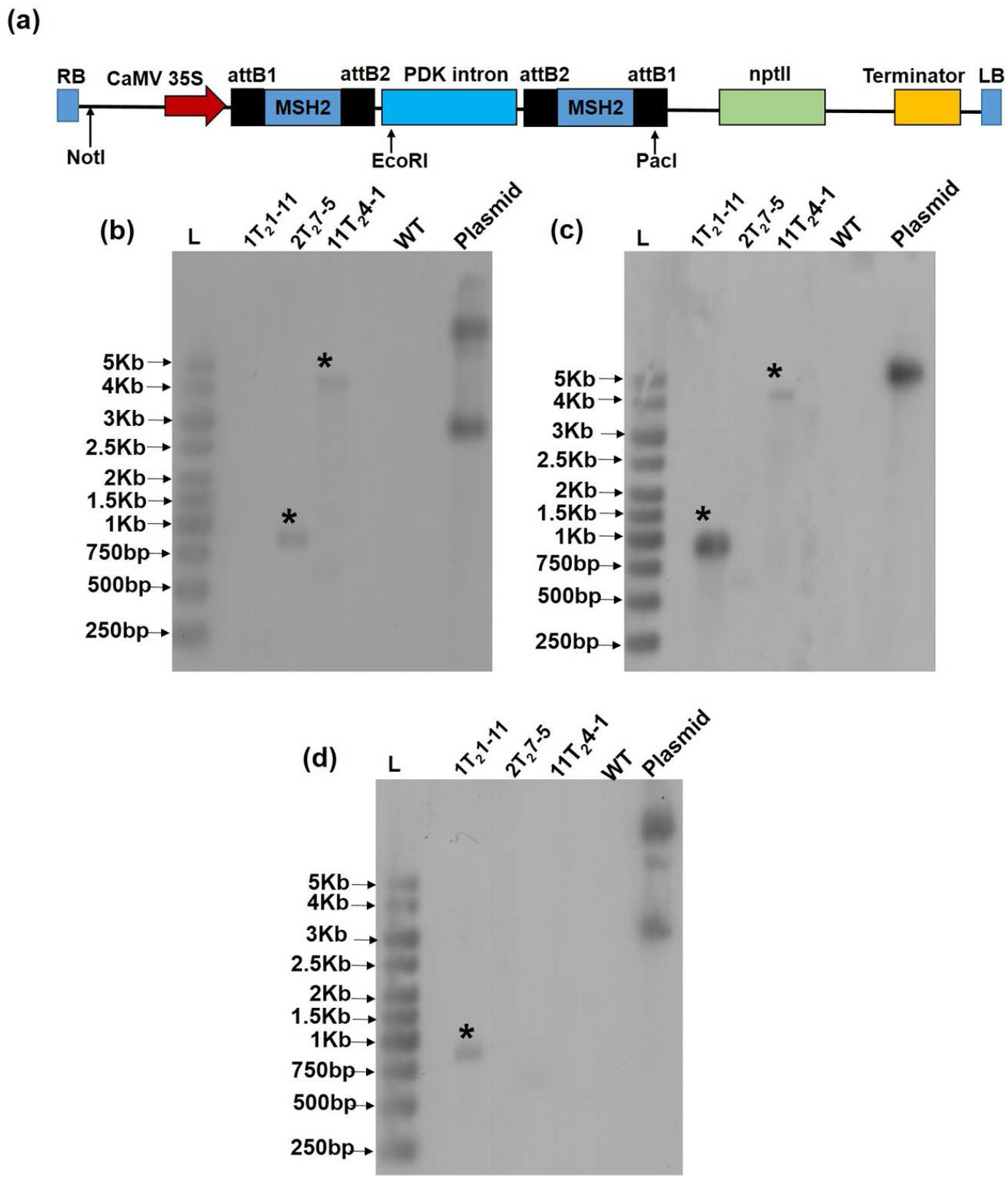
Southern blot of T_2_ *MSH2*-RNAi lines 1T_2_1-11, 2T_2_7-5 and 11T_2_4-1. **(a)** Schematic representation of the construct used for *MSH2* silencing. Black arrow represents the site of the restriction enzyme used for digesting Genomic DNA in Blot **(b)**, **(c)** and **(d)**. DNA was digested with EcoRI **(b)**, with NotI **(c)**, and PacI **(d)**. **L**-1 Kb DNA Ladder (Fermentas). **WT**-Wild-type genomic DNA (negative control); **Plasmid**-, Plasmid DNA bearing *MSH2*-RNAi construct. The blots were probed with radiolabelled *NPTII* sequence. The asterisk (*) indicates the presence of *NPTII* sequence in the transgenic lines.

**Figure S5.**
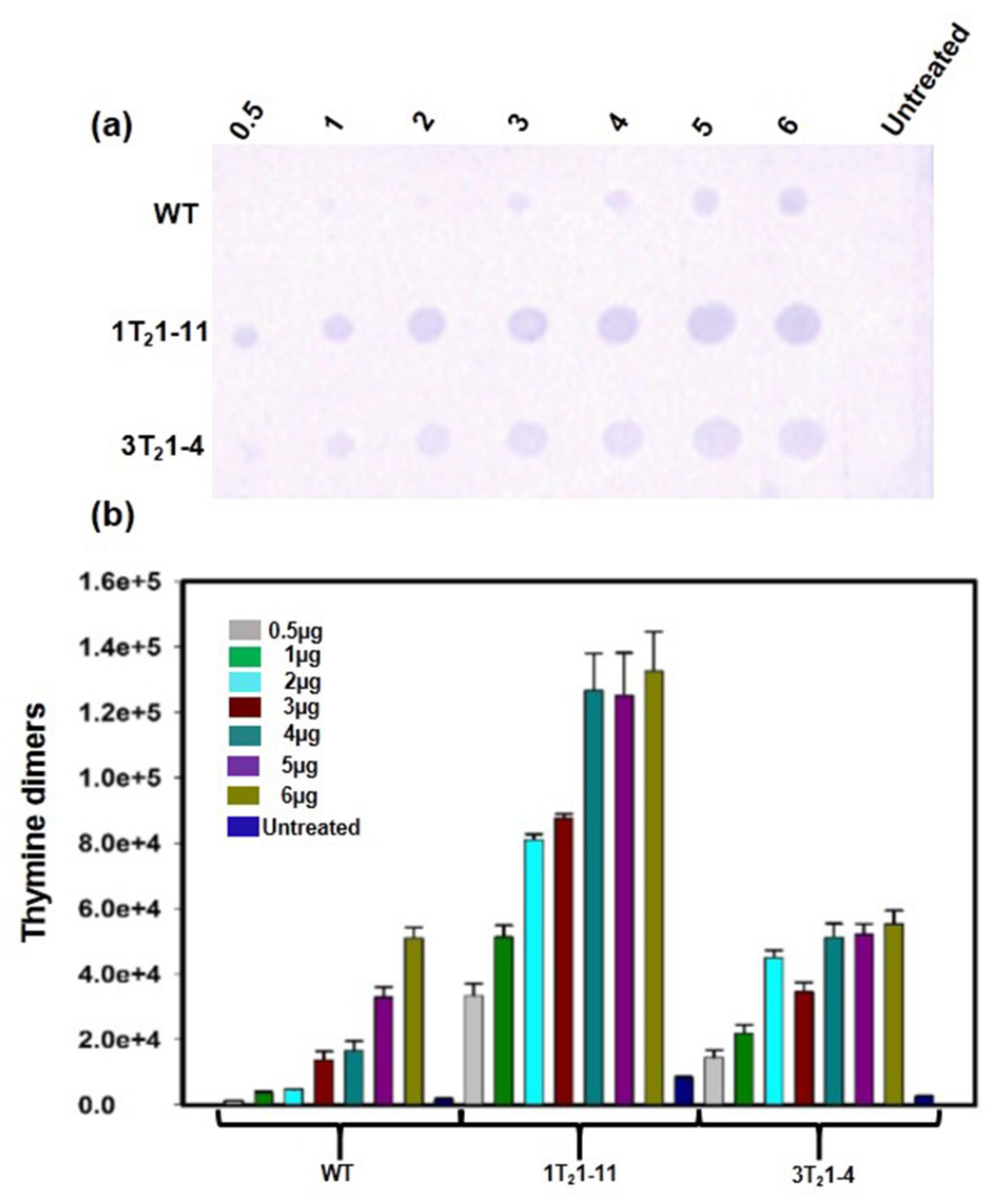
Quantification of thymine dimer levels in genomic DNA of WT and UV-B-treated *MSH2*-RNAi lines, 3T_2_1-4 and 1T_2_1-11. **(a)** Quantification of thymine dimer with different dilutions of genomic DNA. Numbers on top of the blot indicate the spotted amount of genomic DNA (μg) and on the left indicate plant number. **(b)** Quantification of thymine dimer formation in *MSH2*-RNAi lines in **(a)** by ImageJ analysis.

**Figure S6.**
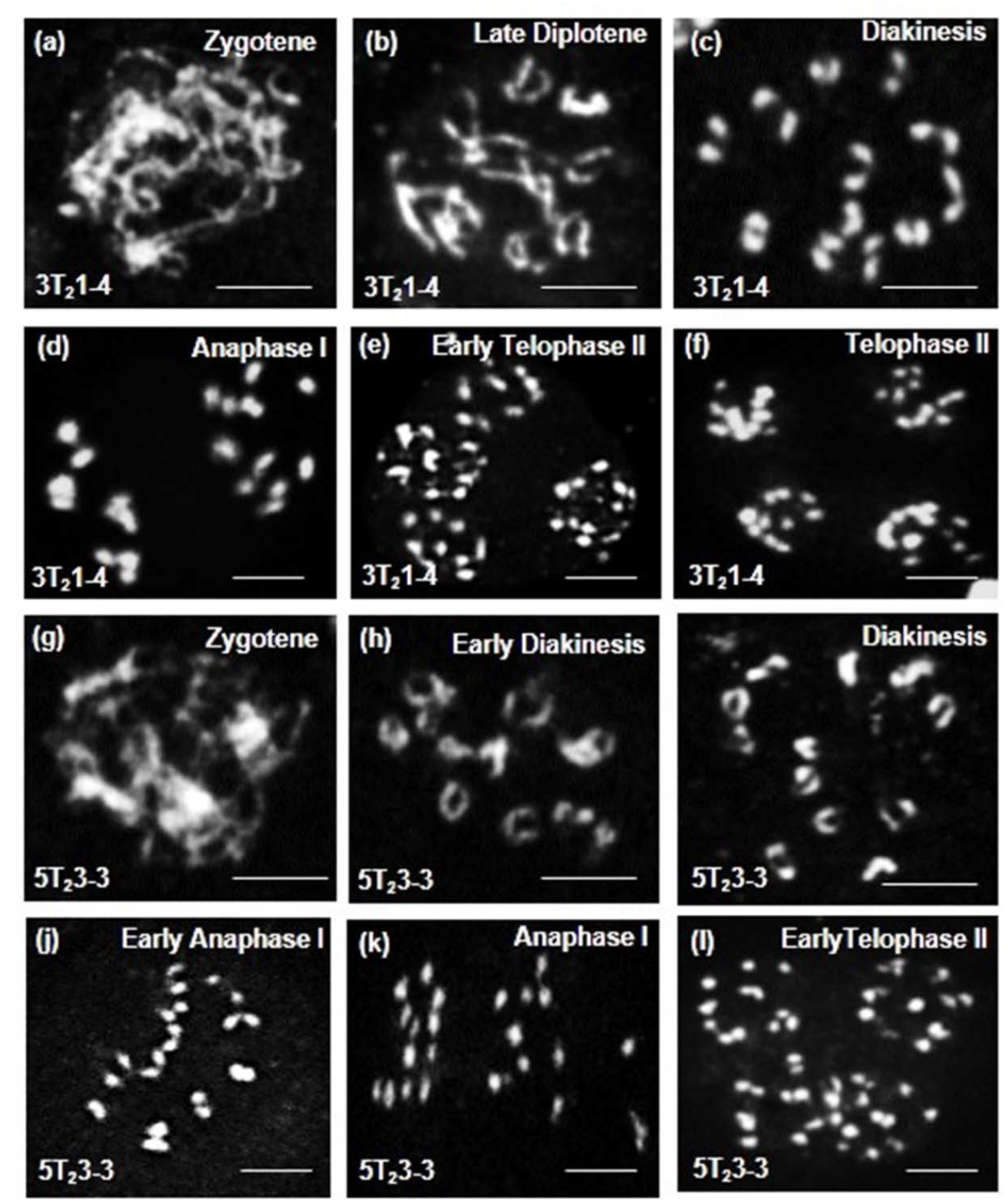
Male meiosis in *MSH2*-RNAi tomato lines. Representative meiotic stages of 3T_2_1-4 **(a-f)** and 5T_2_3-3 **(j-q)** lines from zygotene to telophase. The 3T_2_1-4 line with moderate *MSH2* silencing and 5T_2_3-3 line with no *MSH2* silencing displayed diploid meiotic chromosomes. Scale bar, 10 μm.

**Figure S7.**
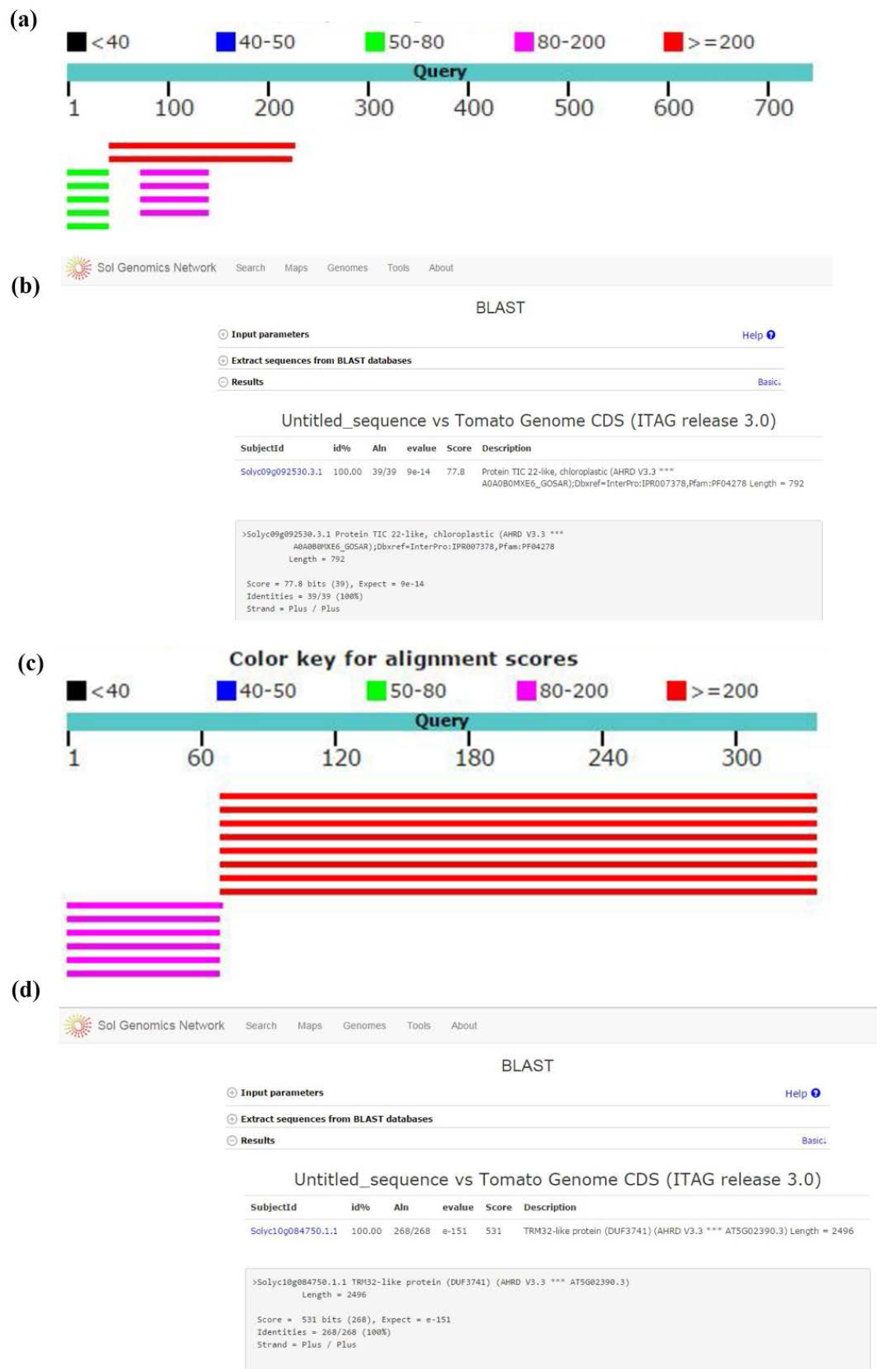
Sequence analysis of FPNI-PCR product of Line No. 1T_2_1-11 and 2T_2_5-5. The FPNI-PCR product from Line 1T_2_1-11 showed 98% homology to tomato chromosome 9 in NCBI gene database **(a)** to *Tic22* gene of tomato in SGN database **(b)**. The FPNI-PCR product from Line 2T_2_ 5-5 showed 100% homology in chromosome 10 in NCBI database **(c)** and to *TRM32* gene of tomato in SGN database **(d)**.

**Figure S8.**
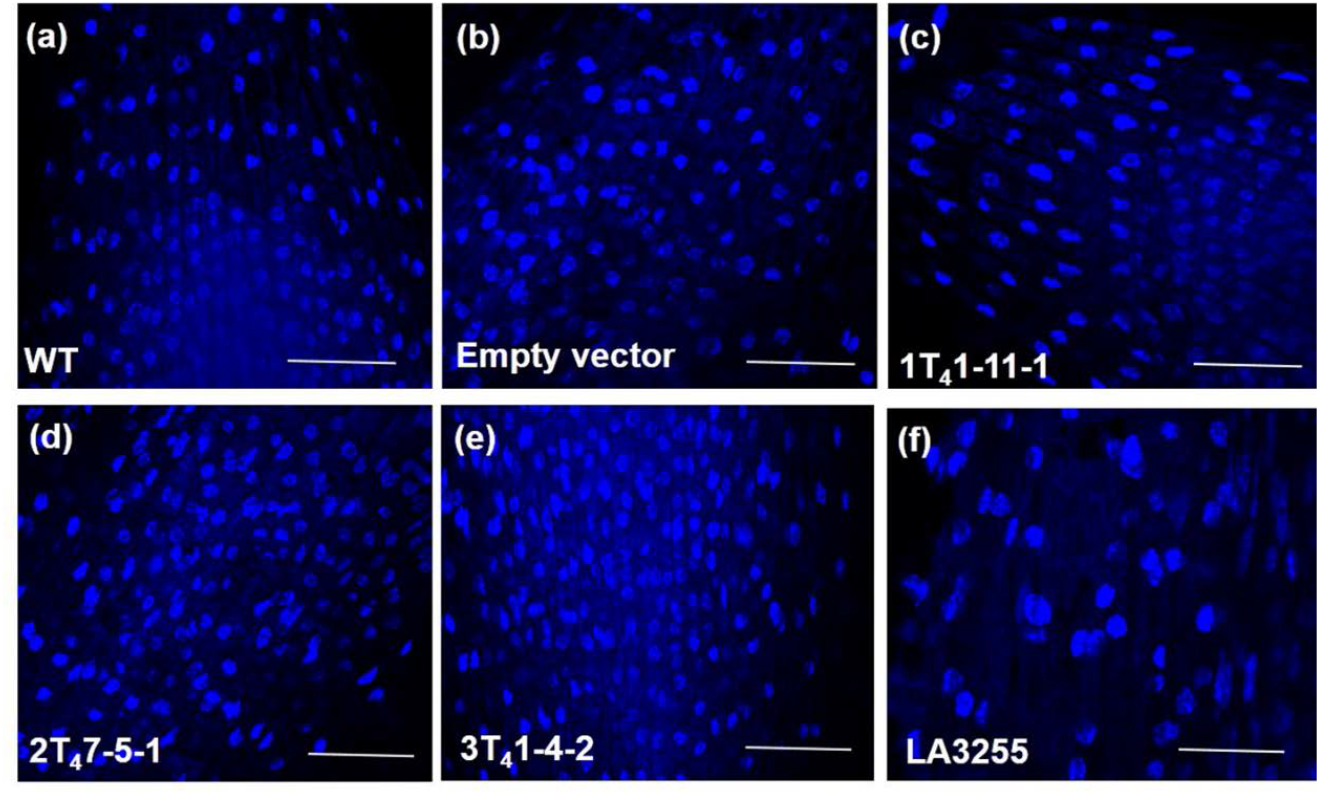
DAPI staining of root tip nuclei of WT and different *MSH2*-RNAi lines. **(a)** WT, **(b)** empty vector control, **(c-e)** progenies of T_3_ *MSH2*-RNAi lines-1T_2_1-11-1 **(c)**, 2T_2_7-5-1 **(d)**, 3T_2_ −4-2 **(e)** and LA3255 **(f)**. Tetraploid tomato line LA3255 shows distinct enlarged tetraploid root tip nuclei in comparison to normal diploid nuclei in WT, control and *MSH2*-RNAi lines. Scale Bar, 50 μm.

**Supplementary Table S1:**
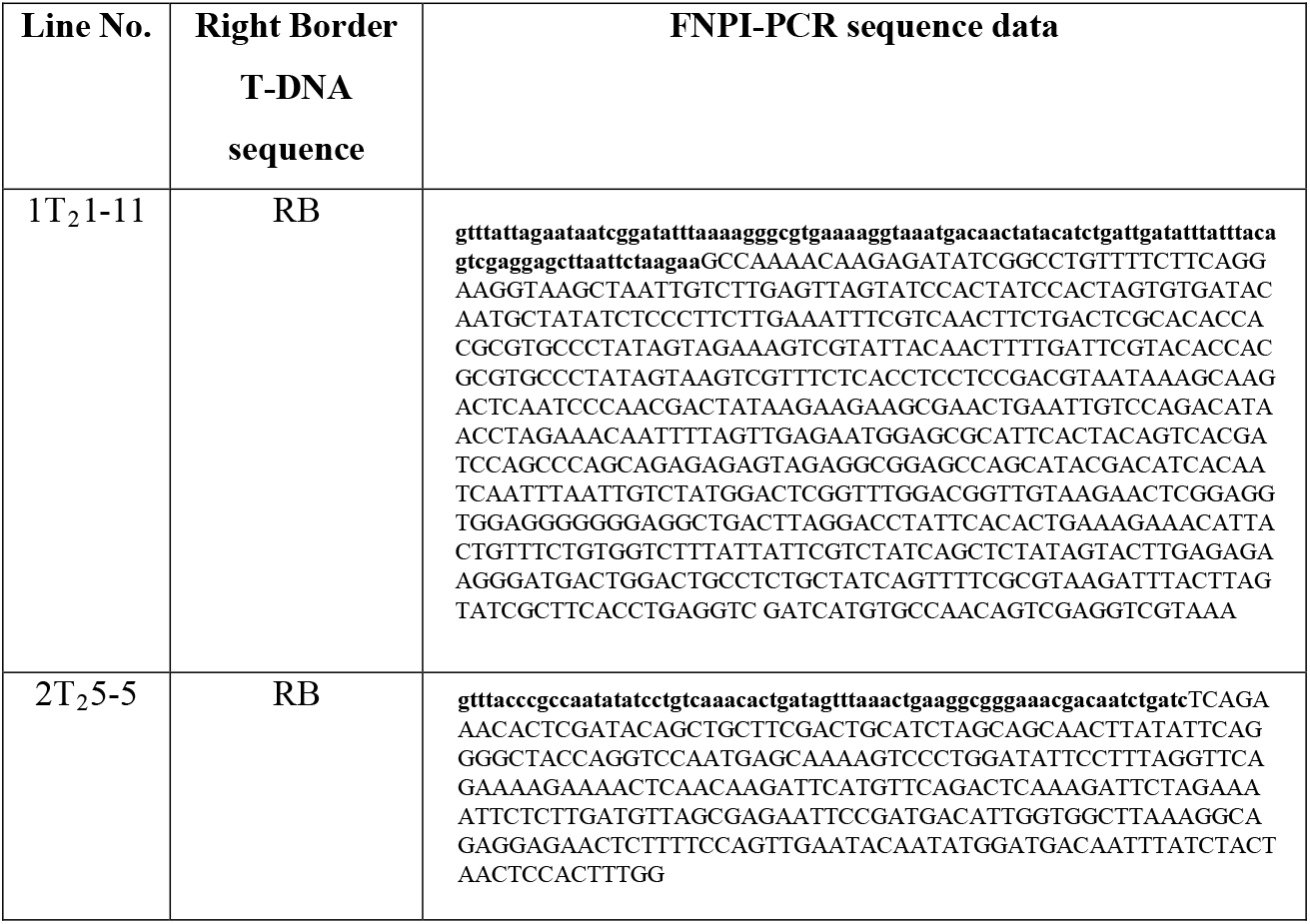
The gene sequences flanking right border of T-DNA obtained using FPNI-PCR. Bold lowercase letters indicate bacterial right border T-DNA sequence and uppercase letters indicate flanking genome sequence from transgenic *MSH2*-RNAi plants.

**Supplementary Table S2.**
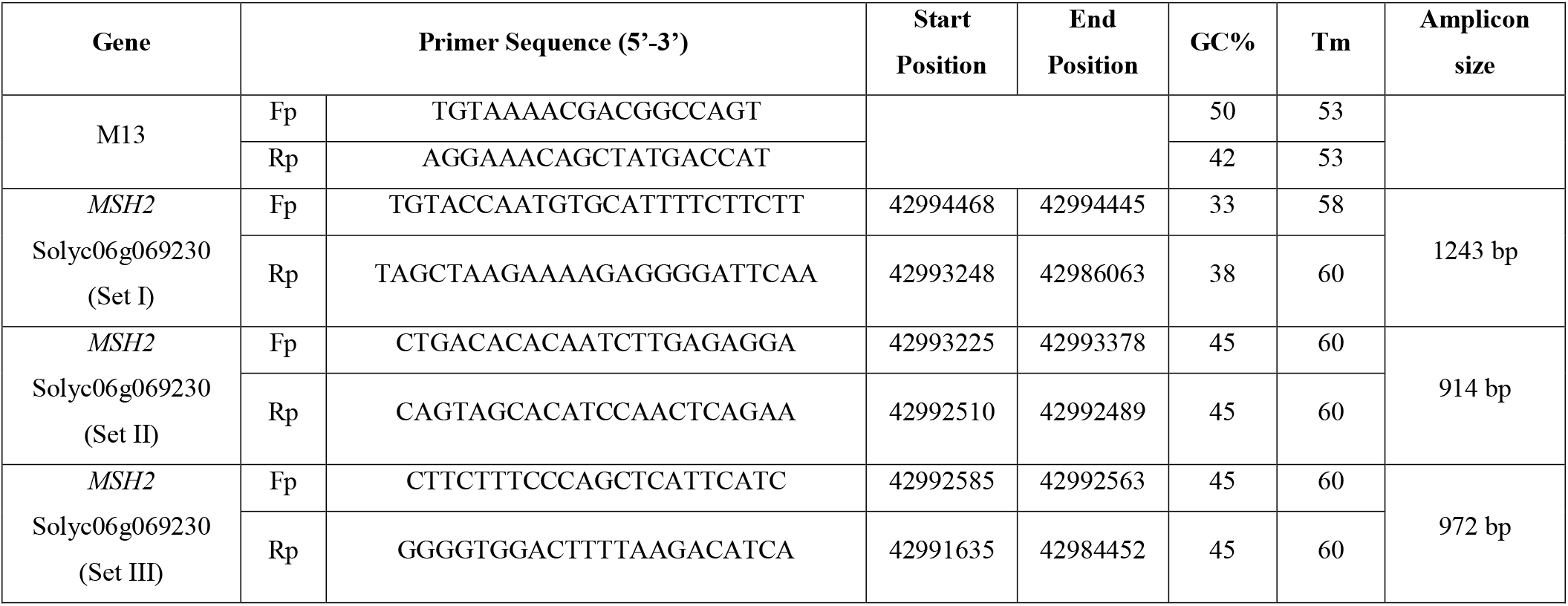
List of genes and the primers used for screening for mutations and SNPs for the regions predicted by CODDLE based on genome sequence of Solanaceae Genome version 2.5. The *MSH2* gene information and location of the primers on genome sequence is also indicated.

**Supplementary Table S3:**
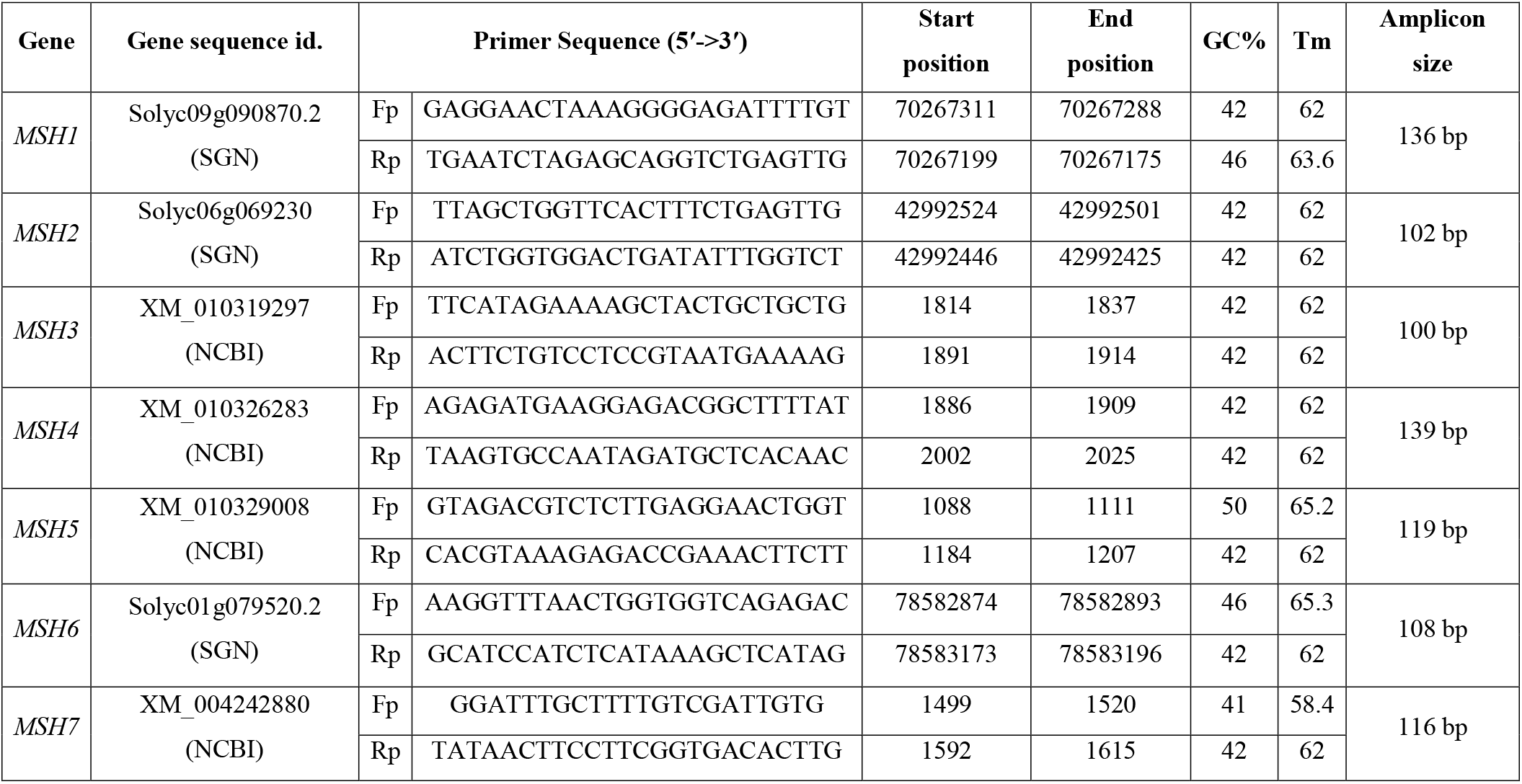
List of genes and the primers used for transcript analysis of MMR pathway.

**Supplementary Table S4:**
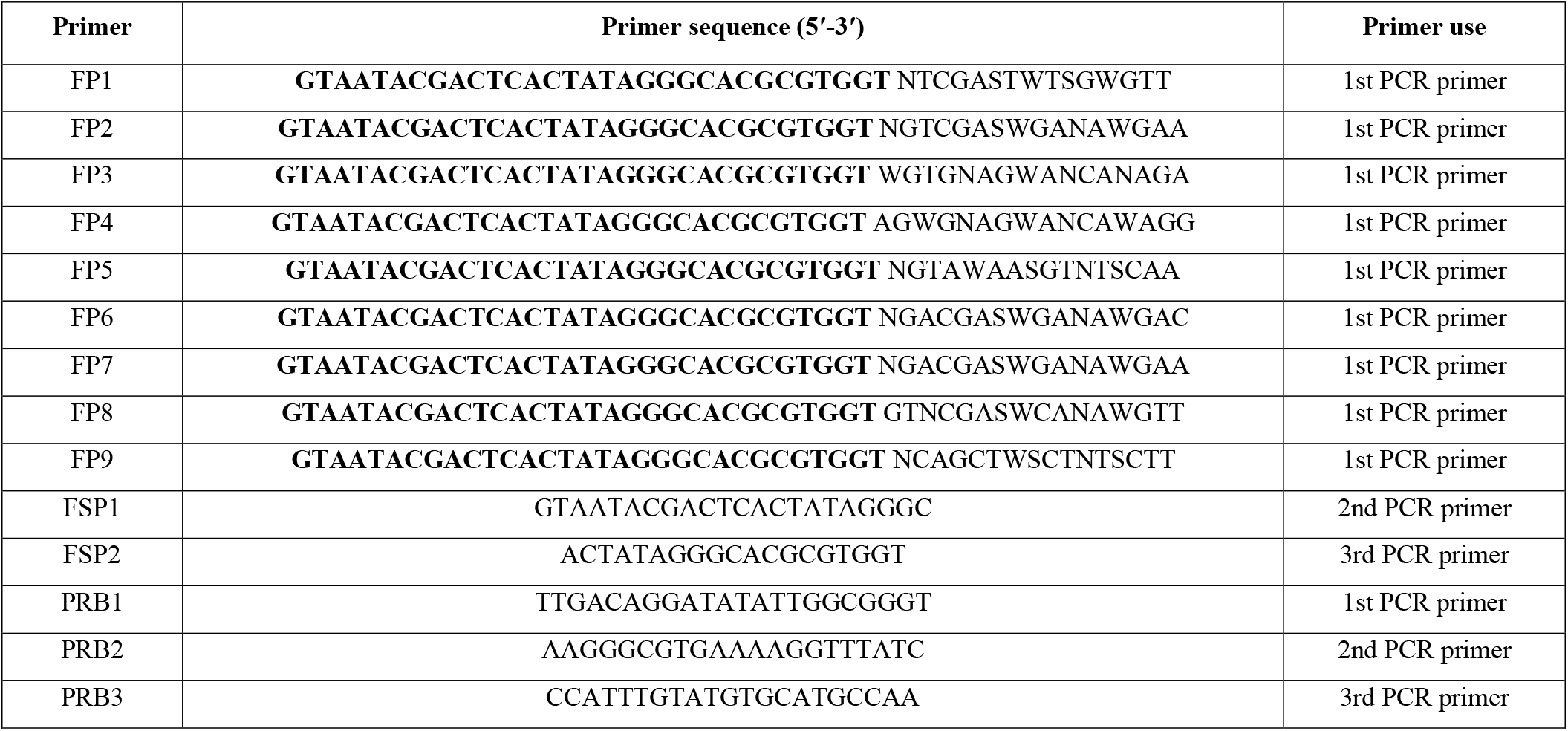
List of genes and the primers used for FNPI-PCR. The common region is highlighted in bold.

